# Acute cerebellar knockdown of *Sgce* reproduces salient features of Myoclonus-dystonia (DYT11) in mice

**DOI:** 10.1101/781005

**Authors:** Samantha G. Washburn, Rachel Fremont, M. Camila Moreno, Chantal Angueyra, Kamran Khodakhah

**Author notes:** Corresponding author: Kamran Khodakhah.

## Abstract

Myoclonus dystonia (DYT11) is a movement disorder caused by loss-of-function mutations in *SGCE* and characterized by involuntary jerking and dystonia that frequently improve after drinking alcohol. Existing transgenic mouse models of DYT11 exhibit only mild motor symptoms, possibly due to rodent-specific developmental compensation mechanisms, which have limited the study of neural mechanisms underlying DYT11. To circumvent potential compensation, we used short hairpin RNA (shRNA) to acutely knock down S*gce* in the adult mouse and found that this approach produced dystonia and repetitive, myoclonic-like movements in mice that improved after administration of ethanol. Acute knockdown of *Sgce* in the cerebellum, but not the basal ganglia, produced motor symptoms, likely due to aberrant cerebellar activity. The acute knockdown model described here reproduces the salient features of DYT11 and provides a platform to study the mechanisms underlying symptoms of the disorder, and to explore potential therapeutic options.

## Introduction

Dystonia is a hyperkinetic movement disorder characterized by involuntary co-contraction of agonist and antagonist muscles resulting in abnormal, repetitive movements and pulled or twisted postures that cause varying degrees of disability and pain (Albanese et al., 2013). The prevalence of dystonia in the general population has been notoriously difficult to determine precisely, due to different methodologies for case classification, but a recent meta-analysis estimated that primary dystonia occurs at a rate of approximately 16 per 100,000 (Steeves et al., 2012). This is likely an underestimate, as cases frequently go undiagnosed and dystonia can also be a secondary symptom in other motor disorders. While dystonia most often occurs sporadically, mutations in several genes have been identified and linked with specific forms of dystonia (Charlesworth et al., 2013). Myoclonus-dystonia (DYT11) is a dominantly inherited form of dystonia caused by loss-of-function mutations in the *SGCE* gene, which encodes the protein epsilon sarcoglycan (ε-SG) (Zimprich et al., 2001). The vast majority of DYT11 patients with *SGCE* mutations inherit the disorder from the male parent, with markedly reduced penetrance in the offspring of female patients (Asmus et al., 2002; Zimprich et al., 2001). The disorder is maternally inherited in only 5-10% of cases. This pattern of dominant, primarily paternal inheritance is consistent with imprinting, or epigenetic silencing, of the maternal allele. Indeed, in DNA and RNA samples from human blood *SGCE* is maternally imprinted, but the imprinting pattern in the brain is unknown (Grabowski et al., 2003).

DYT11 is characterized primarily by involuntary, tic-like jerking of the arms, neck, and trunk (myoclonus) and oftentimes mild to moderate dystonia that worsens with stress (Kyllerman et al., 1990). Motor symptoms usually appear in childhood or adolescence (Nardocci et al., 2008; Raymond and Ozelius, 1993; Raymond et al., 2008) and are frequently accompanied by psychiatric symptoms, including anxiety, panic attacks, and obsessive compulsive disorder (Peall et al., 2011; Weissbach et al., 2013). DYT11 is known as alcohol-responsive dystonia given that approximately half of patients with DYT11 report improvement of motor symptoms after consuming alcohol. Strikingly, for these patients, alcohol tends to provide more therapeutic relief than any available pharmacological interventions, which have frequently been ineffective or poorly-tolerated. Unfortunately, the addictive and neurodegenerative consequences of chronic alcohol use preclude its use as a therapeutic option. Although patients generally live an active life of normal span, the motor and psychiatric symptoms of this disorder can cause a great deal of physical and psychosocial distress, significantly impacting the patients’ quality of life.

A limiting factor in the development of more effective treatment strategies for patients with DYT11 is the lack of understanding of the neural basis of the disorder. While the genetic cause of DYT11 has been known for 10 years, the function of ε-SG protein, particularly in the nervous system, and how loss-of-function of ε-SG leads to motor symptoms remains elusive. This issue is not specific to DYT11, but rather a common problem in understanding genetic dystonias. While mutations in the genes encoding α-, β-, γ-, and δ-sarcoglycans cause different forms of muscular dystrophies, no signs or symptoms of muscle disease have been detected in DYT11 patients with mutations in *SGCE* (Hjermind et al., 2008). Moreover, electrophysiological and fMRI studies have cited a subcortical origin of the disorder (Marelli et al., 2008; Nitschke et al., 2006; Roze et al., 2008). Generally, dystonia has been viewed as a disorder of the basal ganglia. Indeed, in patients with DYT11, changes have been observed in the internal global pallidus (GPi), an output structure of the basal ganglia. In particular, studies have shown that gray matter volume of the GPi (Beukers et al., 2011) and firing patterns of individual neurons in the GPi (Welter et al., 2015) correlates with the severity of motor symptoms in patients. Moreover, deep brain stimulation of the GPi is effective for most DYT11 patients, particularly those with *SGCE* mutations (Azoulay-Zyss et al., 2011; Fernandez-Pajarin et al., 2016; Gruber et al., 2010; Kosutzka et al., 2018; Rocha et al., 2016; Rughani and Lozano, 2013).

On the other hand, recent studies have shown that dystonia is associated with changes in the activity, structure, and connections of the cerebellum (Draganski et al., 2003; Dresel et al., 2014; Hendrix and Vitek, 2012; Jinnah and Hess, 2006; LeDoux et al., 1993; LeDoux et al., 1995; Prell et al., 2013; Shakkottai, 2014; Song et al., 2014). In DYT11 patients, there is evidence from fMRI (Beukers et al., 2010; Nitschke et al., 2006; van der Salm et al., 2013), and PET (Carbon et al., 2013) for increased cerebellar activity. There are also diffusion tensor imaging data to suggest that changes occur in the white matter tracts that connect the cerebellum to the thalamus and other parts of the brain (van der Meer et al., 2012). Furthermore, studies on the expression of *SGCE* and in the human brain and *Sgce*, the mouse homolog of *SGCE*, in the rodent brain have suggested that a brain-specific isoform, which might be responsible for the purely neurophysiological phenotype of DYT11, is highly expressed in the principle computational and output neurons of the cerebellum, the Purkinje cells and deep cerebellar nuclei (DCN) neurons, respectively, while it is expressed at lower levels in the different nuclei of the basal ganglia (Ritz et al., 2011; Xiao et al., 2017; Yokoi et al., 2005). Lastly, the amelioration of symptoms after ethanol ingestion reported by a number of patients with DYT11 points to a role for the cerebellum in this disorder, as the cerebellum is exquisitely sensitive to alcohol. Several labs have demonstrated that alcohol modulates Purkinje cell spiking activity (Chu, 1983; Rogers et al., 1980; Sinclair et al., 1980), increases inhibitory synaptic transmission in the cerebellum (Carta et al., 2004), and alters synaptic plasticity at synapses onto Purkinje cells (Belmeguenai et al., 2008; He et al., 2013).

The goal of this study was to establish a mouse model that would enable the investigation of the neural correlates of motor symptoms in DYT11. Previous genetic knockout models have had subtle effects on motor behavior. Mice with a ubiquitous genetic knockout of S*gce* exhibited full body jerks and had difficulty learning in a balance beam task (Yokoi et al., 2006). Conditional knockout of *Sgce* in Purkinje cells of the cerebellum (Yokoi et al., 2012a) or specifically in the striatum (Yokoi et al., 2012b), the input nucleus of the basal ganglia, produced milder effects. Importantly, none of these mice exhibited dystonia or alcohol-responsiveness, both prominent features of DYT11. Unfortunately, genetic models of hereditary dystonia have often failed to recapitulate many of the key motor symptoms of the disorder. One possibility for this phenomenon is compensation for the gene that has been knocked out during brain development in rodents (Fremont and Khodakhah, 2012).

In recent years, our laboratory has successfully generated symptomatic mouse models of dystonia using an acute knockdown approach to overcome potential developmental compensation (Calderon et al., 2011; Fremont et al., 2014; Fremont et al., 2017; Fremont et al., 2015). Here, we used short hairpin ribonucleic acid (shRNA) to knockdown *Scge* mRNA selectively in the cerebellum or basal ganglia of adult mice. We found that knockdown of *Sgce* in the cerebellum, but not the basal ganglia, produced overt motor symptoms, including dystonia. Consistent with what is seen in patients, administration of ethanol improved motor symptoms in S*gce* knockdown mice but had no effect on motor symptoms of mice with knockdown of the gene that causes primary torsion dystonia (DYT1). We further found that, compared to mice injected with a control shRNA, both Purkinje cells and DCN neurons of the cerebellum fire aberrantly in *Sgce* knockdown mice.

## Results

We hypothesized that developmental compensation for genetic knockout of *Sgce* could account for the mild motor symptoms observed in previous mouse models of DYT11. To address this issue, we employed an acute, shRNA-mediated knockdown strategy to generate a symptomatic mouse model of DYT11. This strategy was successful in two previous studies of rapid onset dystonia-parkinsonism (DYT12) (Fremont et al., 2015) and primary torsion dystonia (DYT1) (Fremont and Khodakhah, 2012; Fremont et al., 2017).

A wealth of evidence suggests that the cerebellum may be particularly important in the pathophysiology of DYT11 (Beukers et al., 2010; Carbon et al., 2013; Ritz et al., 2011; van der Meer et al., 2012; van der Salm et al., 2013). To examine the contribution of the cerebellum to the symptoms observed in DYT11 patients, we injected AAV9 virus encoding shRNA against *Sgce* mRNA and a GFP reporter (AAV-SGCEshRNA-GFP) into the cerebellum of adult mice (Figure 1A). We found that injection of AAV-SGCEshRNA-GFP resulted in expression of the construct throughout the cerebellum without causing any gross morphological abnormalities (Figure 1B), as well as significant knockdown of *Sgce* mRNA as determined by qRT-PCR (Figure 1C, Mann-Whitney Test, WT vs. NT CB: p = 0.7922; NT CB vs. sgce KD CB 1: p = 0.0079; NT CB vs. sgce KD CB 2: p = 0.0159; N_WT_ = 6, N_NT CB_ = 5, N_sgce KD CB 1_ = 5, N_sgce KD CB 2_= 4). To control for shRNA-specific effects, a group of mice were injected with shRNA that does not target any gene in the genome (AAV-shNT-GFP, NT CB). To control for off-target effects of the shRNA, we used two sequences of shRNA that target separate regions of *Sgce* (*Sgce* KD CB 1 and *Sgce* KD CB 2). Mice were injected with either of two shRNA sequences against *Sgce* into the cerebellum. Their behavior in the open field was assessed by four scorers blind to the condition of the animal using a previously published dystonia scale, where a score greater than or equal to 2 indicates dystonia (Calderon et al., 2011; Fremont et al., 2014; Fremont et al., 2017; Fremont et al., 2015). Sgce KD CB mice have a time-dependent induction of dystonia, indicated by a dystonia score of 2 or above (Video 1, Figure 1Di, sgce KD CB 1: N = 39; sgce KD CB 2: N = 40). Abnormal movements and postures were observed throughout the body of the animal, although the most severe dystonic postures were observed in the hind limbs and tails (Figure 1D, ii). In contrast, mice injected with AAV-shNT-GFP did not develop symptoms (Video1, Figure 1Di, NT CB: N = 16). The development of dystonic symptoms in *Sgce* KD CB mice is time-dependent because the transfection of the virus, expression of the shRNA, and knockdown of ε-SG protein increases over time. This provides the unique opportunity for within-animal control to make sure that the surgery itself did not cause motor symptoms. Accordingly, the dystonia scores for sgce KD CB 1 and sgce KD CB 2 mice for time points of 2 weeks or more after injection were significantly different from the dystonia scores of the same animals at < 1 week (Figure 1Di, p < 0.01, Wilcoxon matched-pairs signed rank test). Consistent with these data, the dystonia scores of sgce KD CB 1 and sgce KD CB 2 mice at < 1 week after injection were not significantly different from NT CB mice at the same time point (Figure 1Di, p = 0.81 and p = 0.97, respectively, t-test, Holm-Sidak method). The dystonia scores of sgce KD CB 1 and sgce KD CB 2 were not significantly different (P> 0.05, t-test, Holm-Sidak method).

**Figure 1.**
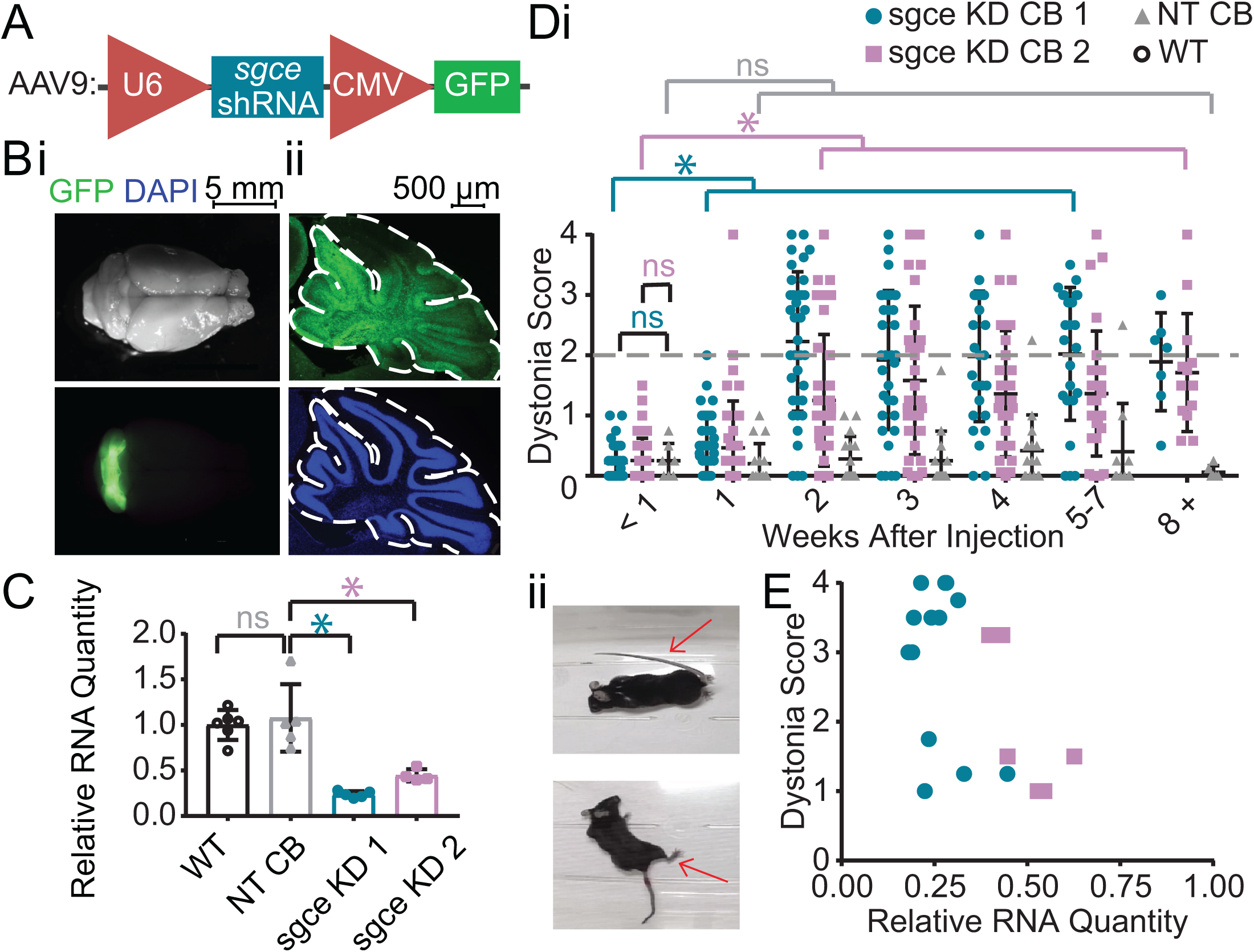
shRNA-mediated knockdown of Sgce in the cerebellum causes dystonia. (A) Schematic of AAV-shSGCE-GFP construct. (B) Images of the whole brain (i) and sagittal cerebellar section (ii) from an sgce KD CB mouse. (C) Quantification of qRT-PCR confirms that *Sgce* RNA is reduced *in vivo*. (Mann-Whitney Test, WT vs. NT CB: p = 0.7922; NT CB vs. sgce KD CB 1: p = 0.0079; NT CB vs. sgce KD CB 2: p = 0.0159; NWT = 6, NNT CB = 5, Nsgce KD CB 1 = 5, Nsgce KD CB 2= 4). (D,i) Injection of sgce KD 1 and 2 into the cerebellum produced dystonia, while injection of NT did not. (sgce KD CB 1: N = 39; sgce KD CB 2: N = 40; NT CB: N = 16). Dystonia was measured on a previously published dystonia scale by four scorers blind to the condition of the animal. A score greater than or equal to 2 indicates dystonia. The dystonia scores for sgce KD CB 1 and sgce KD CB 2 mice for time points of 2 weeks or more after injection were significantly different from the dystonia scores of the same animals at < 1 week (Wilcoxon matched-pairs signed rank test, p < 0.01). The dystonia scores of sgce KD CB 1 and sgce KD CB 2 mice at < 1 week after injection were not significantly different from NT CB mice at the same time point (t-test, Holm-Sidak method, p = 0.81 and p = 0.97, respectively). (ii) Example dystonic postures exhibited by sgce KD CB mice. (E) Scatter plot of RNA levels normalized to the mean of WT, determined by qRT-PCR, plotted against the Dystonia Score observed in a subset of animals injected with varying concentrations of shRNA (WT: N = 5, NT: N = 5, sgce KD 1: N = 13, sgce KD 2: N = 7).

Approximately 75% (29/39) of sgce KD CB 1 mice and 60% (23/40) of and sgce KD CB 2 mice developed dystonia. To explore whether the absence of symptoms is due to inefficient knockdown of *Sgce* or whether it is due to penetrance of the phenotype, we plotted the extent of *Sgce* knockdown, determined by qRT-PCR, against the dystonia score recorded in a subset of animals (Figure 1E). Based on these data, there appears to be a correlation between the severity of symptoms and knockdown of *Sgce* RNA, consistent with previous data in an shRNA-mediated mouse model of DYT1 (Fremont et al., 2017). However, we found that while 5/20 *Sgce* KD CB animals had more than 50% knockdown of *Sgce* RNA compared to WT, they had a dystonia score of less than 2. It is thus possible that, while the extent of *Sgce* knockdown explains some variability observed in the behavioral data, incomplete penetrance of the phenotype may also contribute to the variability of the symptoms observed in *Sgce* KD CB mice.

Upon closer examination of sgce KD CB mice, we noticed that in addition to dystonia, these mice exhibited a range of motor symptoms that we had not observed in some of the other mouse models of movement disorders previously characterized in our laboratory (Video 2). Sgce KD CB mice had difficulty ambulating normally and showed unsteady or mildly ataxic gate, and myoclonic-like movements in addition to overt dystonia. Furthermore, when suspended by the tail, sgce KD CB mice would spin vigorously and, often, continuously. In order to capture the range of motor symptoms observed in sgce KD CB mice more accurately, we generated two new scales to measure the characteristic symptoms seen in this model. The Disability Scale (Table 1) took into consideration the frequency of sustained dystonic-like postures and repetitive movements, as well as the level of motor impairment. The Spinning Scale (Table 2) was generated to measure the frequency and duration of spinning during a tail suspension test, a symptom unique to sgce KD CB mice compared to our other models of dystonia. The average Spinning Score for a subset of sgce KD CB 1 and sgce KD CB 2 was 2.22 ± 0.77 and 2.00 ± 0.88, respectively (Mean ± S.D., N = 13 and 7). The average Disability Score was 3.02 ± 0.76 and 2.89 ± 0.79 (Mean ± S.D., N = 14 and 8), respectively.

**Table 1.** Disability scale for assessing motor impairment in *Sgce* knockdown mouse model of DYT11.

|  |  |
| --- | --- |
| 0 | **Normal** -- animal has no observable motor deficit |
| 1 | **Slight motor disability** -- animal may exhibit unsteady gait or uncoordinated movement, but does not have any repetitive movements or sustained postures |
| 2 | **Mild motor disability** -- ambulation is mildly impaired; animal exhibits wide stance, very unsteady gait, back-walking, and/or occasional repetitive movements or sustained postures |
| 3 | **Moderate motor disability** -- ambulation is moderately impaired; animal does not properly ambulate, dragging itself on its side, and may exhibit frequent sustained dystonic-like postures or repetitive movements |
| 4 | **Severe motor disability** -- animal is unable to properly ambulate for most of the duration of the observation, exhibits frequent repetitive movements, and/or sustained dystonic-like postures |

**Table 2.** Spinning scale for assessing abnormal motor behavior in *Sgce* knockdown mouse model of DYT11

|  |  |
| --- | --- |
| 0 | **Not present** -- Animal does not spin |
| 1 | **Mild** -- Animal spins intermittently or clearly struggles more than the wild-type |
| 2 | **Moderate** -- Animal spins moderately, making several successive complete rotations |
| 3 | **Severe** -- Animal spins vigorously for most of the duration of the observation |

To examine the contribution of the basal ganglia to symptoms in DYT11, we injected shRNA into the striatum and GPi of wild-type mice (sgce KD BG 1 and 2, Figure 2A). Mice injected with AAV-SGCEshRNA-GFP into the basal ganglia developed mildly abnormal motor symptoms, but they did not develop dystonia (Figure 2B, Video 3, sgce KD BG 1: N = 4, sgce KD BG 2: N = 14). In 5 out of 47 observations, animals scored 2 or above on the Dystonia Scale. Unlike the symptoms observed in sgce KD CB mice, which occurred multiple times in the same recording period and persisted over time, these postures did not occur more than once in the same mouse, suggesting that they might have been false positive identification of dystonic-like postures in those incidences. There were no significant differences in the motor symptoms of sgce KD BG 1 and sgce KD BG 2 mice; consequently, only a small subset of mice was injected with sgce KD BG 1 to confirm the observations made in a larger cohort of sgce KD BG 2 mice. Motor abnormalities were not observed in mice injected with AAV-shNT-GFP into the basal ganglia (Figure 2B, Video 3, NT BG, N = 9).

**Figure 2.**
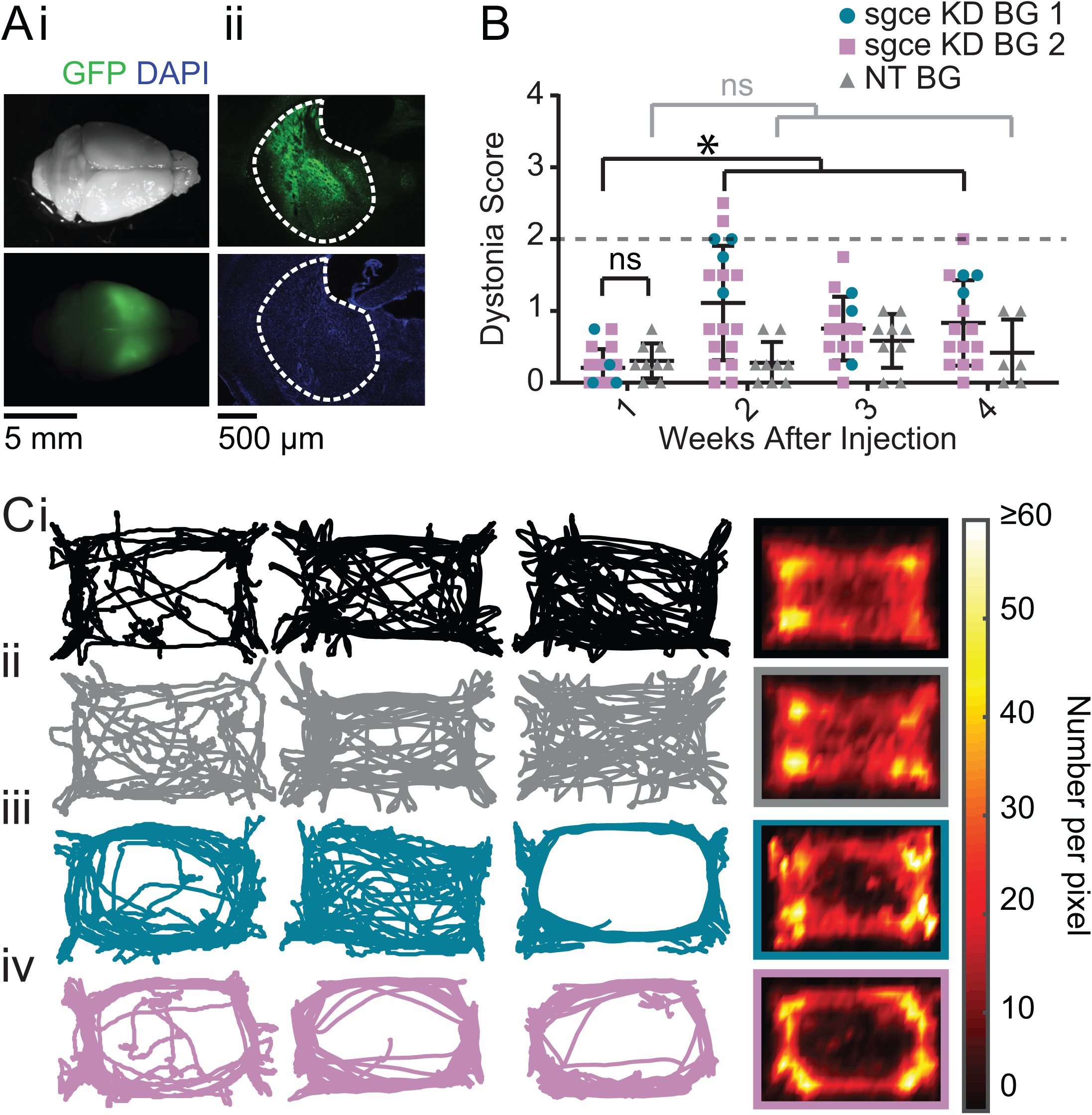
shRNA-mediated knockdown of *Sgce* in the basal ganglia causes motor abnormalities but does not cause overt dystonia. (A) Images of the whole brain (i) and coronal section (ii) from an sgce KD BG mouse. (B) Injection of sgce KD- or NT-shRNA into the basal ganglia did not produce dystonia, as indicated by a score greater than 2 on the Dystonia scale. (sgce KD BG 1: N = 4; sgce KD BG 2: N = 14; NT BG: N = 9). The dystonia scores for sgce KD BG mice for time points of 2 weeks or more after injection were significantly different from the dystonia scores of the same animals at 1 week (Wilcoxon matched-pairs signed rank test, p < 0.001). The dystonia scores of sgce KD BG mice at 1 week after injection were not significantly different from NT BG mice at the same time point (t-test, Holm-Sidak method, p =0.36). (C) sgce KD BG 1 (iii) and sgce KD BG 2 mice (iv) appeared to ambulate more in the periphery of the open field chamber than wild-type (i) and NT BG (ii) mice. The first three columns show example tracks from individual mice. The last column depicts the average, which reflects the number of times the center of mass was detected at a pixel in the arena, and excludes frames where the animal did not move. (WT: N = 12, NT BG: N = 12, sgce KD BG 1: N = 4; sgce KD BG 2: N = 13)

As discussed, although sgce KD BG mice did not develop dystonia, they exhibited abnormal motor activity and these were indicated as scores of 1 and 2 on the Dystonia Scale. These higher scores were statistically significant at the three time points examined. Closer examination of the motor behavior of the mice showed that, in many of them, their activity in the periphery of the open field arena was increased; several Sgce KD BG mice appeared to travel around the periphery and avoid the center of the open field chamber more than either WT or NT BG mice (Figure 2C). In addition to ambulating more on the periphery, these sgce KD BG animals also appeared to spend less time in the center of the open field chamber, compared to the periphery (Figure 2 – Supplement 1A). While the tracks of many of the animals were clearly qualitatively different, there was considerable variability as to how the tracks looked in each animal, and as an average, no statistically significant differences were detected in the ratio of average time spent in the center vs. the periphery (Figure 2 – Figure Supplement 1B). The individual variability from mouse to mouse could be either a function of the extent of knock down of the protein in the BG, or the extent of coverage of the BG by the injected virus. Our ability to detect statistically significant differences in this analysis was also likely limited by the size of the open field chamber used in our experiments. Similarly, total distance traveled was not statistically different among the groups of mice (Figure 2 – Figure Supplement 1C). Sgce KD CB mice did not show the same qualitative preference for the periphery and avoidance of the center observed in sgce KD BG mice (Figure 2 – Supplement 2).

A prominent feature of DYT11 that has not yet been reported in any mouse model is the ability of alcohol to lessen the severity of motor symptoms. To examine whether alcohol could improve the motor symptoms observed in sgce KD CB mice, and further validate this new model, we examined the motor behavior of dystonic (dystonia score ≥ 2) sgce KD CB mice in the open field before and after a subcutaneous injection of ethanol (2 g/kg) or physiological saline. To capture the full range of motor symptoms that ethanol might affect, we scored the motor behavior of these mice on all three scales. Consistent with what has been reported by patients with DYT11, ethanol relieved motor symptoms in sgce KD CB mice for up to 90 minutes after injection as measured on both the Disability Scale (Video 4, Figure 3A, p < 0.0001, 1way ANOVA, N = 16) and Spinning Scale (Figure 3B, p < 0.0001, 1way ANOVA, N = 19). The reduction in Disability and Spinning Scores were significant at each time point after correcting for multiple comparisons (Holm-Sidak’s multiple comparisons test). Similarly, ethanol reduced motor dysfunction as measured on the Dystonia Scale (Figure 3C, p < 0.0001, 1way ANOVA, N = 16). The reduction in dystonia after ethanol was significant at each time point in sgce KD CB mice after correcting for multiple comparisons (Holm-Sidak’s multiple comparisons test). To determine whether this effect was specific to sgce KD CB mice, the effect of alcohol was examined in another symptomatic shRNA-mediated mouse model of primary torsion dystonia (DYT1). Alcohol-responsiveness was not observed in this model of DYT1 (Fremont et al., 2017), although they exhibit overt dystonia, suggesting that the response to alcohol is specific to ε-SG knockdown (Video 4, Figure 3D, 0.2391, 1way ANOVA, N = 5). These findings show that acute knockdown of ε-SG in the cerebellum of adult rodents is sufficient to generate ethanol-responsive motor symptoms, including dystonia.

**Figure 3.**
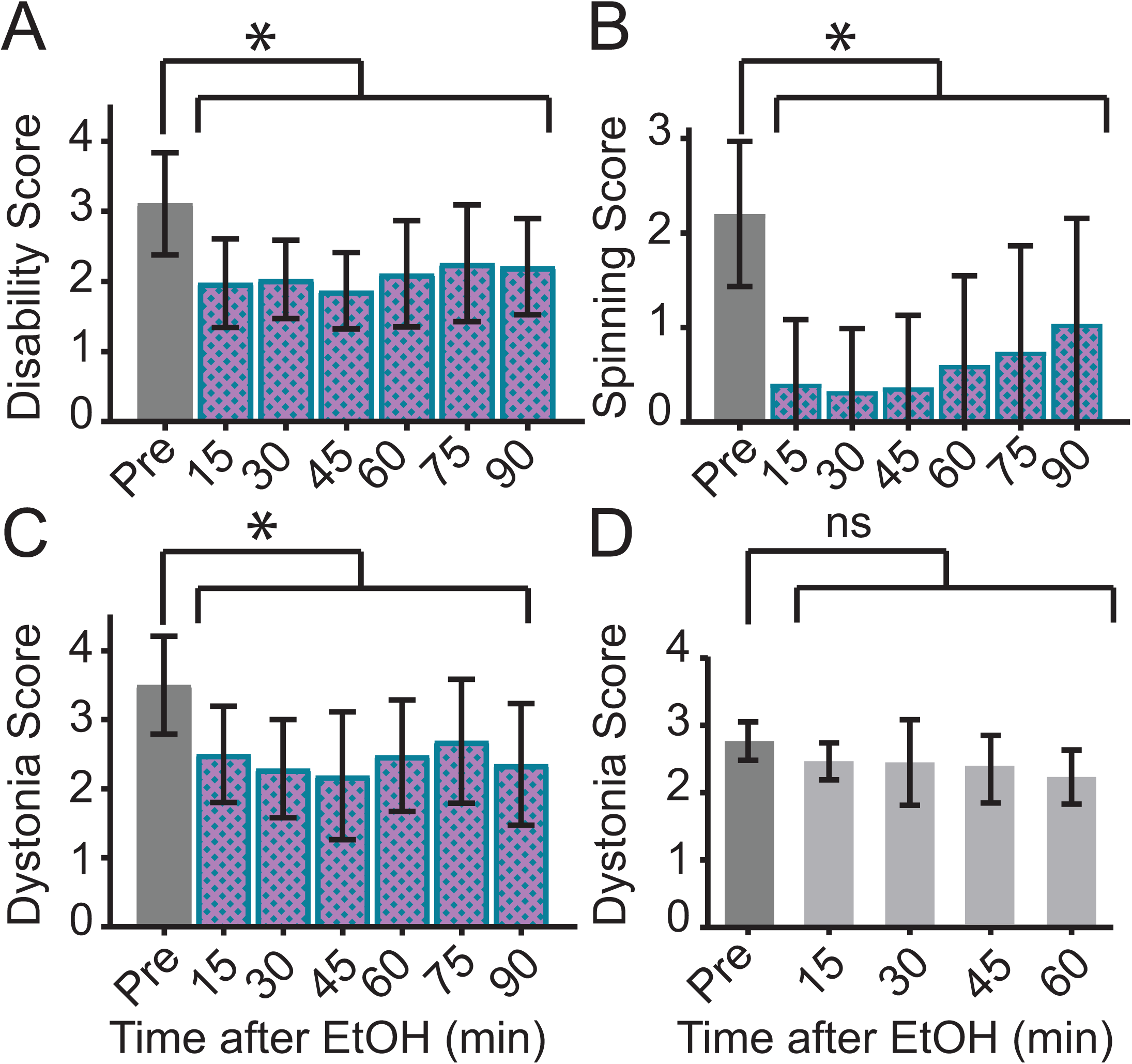
Ethanol relieves motor symptoms in sgce KD CB, but not tor1a KD CB, mice. (A) Disability score of sgce KD CB mice after ethanol injection. Ethanol reduces the disability score of mice injected with shRNA against sgce. Alleviation of symptoms persisted for up to 90 minutes after ethanol (p < 0.0001, 1way ANOVA, Mean + S.D., N = 16). (B) Spinning score of sgce KD CB mice after ethanol injection. Ethanol reduces the spinning score of mice injected with shRNA against sgce. (p < 0.0001, 1way ANOVA, Mean + S.D., N = 19). (C) Dystonia score of sgce KD CB mice after ethanol injection. Ethanol significantly reduced the dystonia score of sgce KD CB mice (p < 0.0001, 1way ANOVA, Mean + S.D., N = 16). (D) Dystonia score of tor1a KD mice after ethanol injection. Ethanol had no effect of the dystonia score, which reflects the primary symptoms caused by tor1a knockdown, in mice injected with shRNA against tor1a (p = 0.2391, 1way ANOVA, Mean + S.D., N = 5).

Although ethanol reduced motor symptoms on the Disability, Spinning, and Dystonia Scales in Sgce KD CB mice, it did not consistently alter the distance traveled in the open field in each mice (Figure 3 – Figure Supplement 1). While, on average, there was a significant increase in the distance traveled in the open field after EtOH (p = 0.0148, 1Way ANOVA), this was driven largely by a small number of mice, and the effect was not statistically significant after correcting for multiple comparisons (Dunnett’s multiple comparisons test, p> 0.05 at each time point). In contrast to EtOH, saline had no effect on the Dystonia Score or distance traveled in sgce KD CB mice (Figure 3 – Supplement 2, Dystonia Score: p = 0.9517, Distance Traveled: p = 0.2851, 1way ANOVA).

Since we successfully generated a symptomatic mouse model of DYT11, we further sought to examine the underlying neural correlates of motor symptoms in DYT11 using electrophysiological techniques. We hypothesized that if irregular activity of the cerebellum contributes to the motor symptoms in sgce KD CB mice, then the output of the cerebellum, the DCN neurons, must be affected. To that end, we performed extracellular single unit recordings from DCN neurons in awake, head-restrained sgce KD CB or NT CB mice. We found that shRNA knockdown of *Sgce* mRNA in the cerebellum caused aberrant activity of cerebellar output neurons (Figure 4). Specifically, we found that the firing rate of DCN neurons in sgce KD CB mice was reduced (NT CB = 62.9 ± 24.8 spikes/s, n = 9, N = 4 and sgce KD CB = 32.2 ± 19.5 spikes/s, n = 32, N = 8, Mean ± S.D.; Welch’s t-test, p = 0.0057). At the same time, the interspike interval coefficient of variation (ISICV) of DCN neurons was increased in sgce KD CB mice (NT CB = 0.50 ± 0.16 and sgce KD CB = 1.00 ± 0.61, Mean ± S.D.; Welch’s t-test, p = 0.0002). The ISICV, a frequently used measure of the regularity of the firing rate of a cell, is the standard deviation of the ISI divided by the mean ISI and thus dependent on the average firing rate of the cell. Sgce KD CB mice had a decreased average firing rate, which would result mathematically in a decreased ISICV if the standard deviation of the ISI was unchanged. The significant increase in the ISI CV in sgce KD CB mice demonstrates that despite the decrease in average firing rate, the cells are firing with much greater variability in their ISIs. There were no significant differences in the mode firing rates of sgce KD CB mice and NT CB mice (NT CB = 80.4 ± 36.6 and sgce KD CB = 74.8 ± 53.5 spikes/s, Mean ± S.D.; Welch’s t-test, p = 0.73). The change in ISICV was primarily a consequence of more pauses in DCN neurons in sgce KD CB mice, rather than a shift to high-frequency burst firing. There was a clear broadening of the curve of the ISI histogram in sgce KD CB mice (Figure 4F), and no increase in the autocorrelation of neurons at time points very close to zero (Figure 4G).

**Figure 4.**
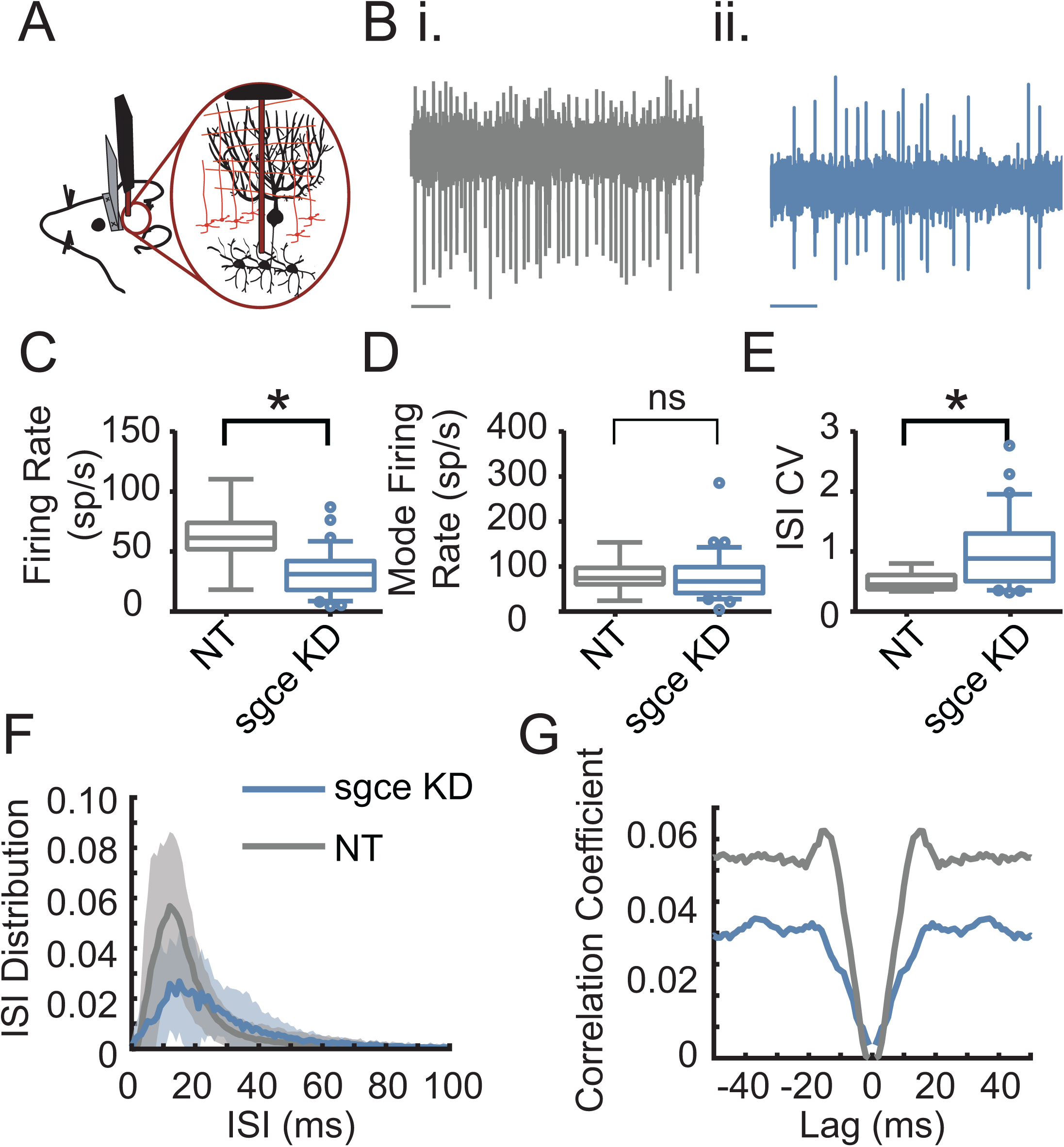
Cerebellar nuclei neurons fire aberrantly in sgce KD CB mice. (A) Experimental schematic. Extracellular electrophysiological recordings were made from neurons in the cerebellar nuclei in awake, head-restrained mice. (B) Example traces from a mouse injected with non-targeting shRNA (i) and shRNA against *Sgce* (ii). Scale bar represents 100 ms. (C) Average firing rates of DCN neurons in NT CB and sgce KD CB animals (NT CB = 62.9 ± 24.8 spikes/s, n = 9, N = 4 and sgce KD CB = 32.2 ± 19.5 spikes/s, n = 32, N = 8, Mean ± S.D.; Welch’s t-test, p = 0.0057). (D) Mode firing rates of DCN neurons in NT CB and sgce KD CB animals (NT CB = 80.4 ± 36.6 and sgce KD CB = 74.8 ± 53.5 spikes/s, Mean ± S.D.; Welch’s t-test, p = 0.73). (E) Interspike interval coefficients of variation of DCN neurons in NT CB and sgce KD CB animals (NT CB = 0.50 ± 0.16 and sgce KD CB = 1.00 ± 0.61, Mean ± S.D.; Welch’s t-test, p = 0.0002). (F) Normalized ISI histogram of DCN neurons in NT CB and sgce KD CB mice. (G) Autocorrelogram of DCN neurons in NT CB and sgce KD CB mice.

The Purkinje cells, the principle cells of the cerebellar cortex and its sole output, provide powerful inhibitory synaptic input onto DCN neurons. The irregular firing activity observed in DCN neurons could partly be due to aberrant Purkinje cell input. To examine this possibility, we performed extracellular recordings from Purkinje cells in awake, head-restrained sgce KD CB and NT CB mice (Figure 5). Similar to what was observed in DCN neurons, sgce KD in the cerebellum reduced the average firing rate of Purkinje cells (NT CB = 53.3 ± 24.1 spikes/s, n = 30, N = 4 and sgce KD CB = 39.2 ± 26.6 spikes/s, n = 57, N = 11, Mean ± S.D.; Welch’s t-test, p = 0.0028), and increased their ISICV (NT CB = 0.60 ± 0.25 and sgce KD CB = 1.06 ± 0.57, Mean ± S.D.; Welch’s t-test, p < 0.0001). In contrast to DCN neurons, Purkinje cells in sgce KD CB mice had an increased mode firing rate (NT CB = 77.5 ± 41.6 spikes/s and sgce KD CB = 115.0 ± 99.4 spikes/s, Mean ± S.D.; Welch’s t-test, p = 0.0158). This is consistent in the rightward shift of the curve in the ISI histogram (Figure 5F) and the observation of longer pauses in DCN neurons. Like DCN neurons, Purkinje cells did not show an increase in the autocorrelation of neurons at time points very close to zero (Figure 5G), suggesting that the neurons are not necessarily burst firing at high frequencies, despite the increase in the predominant firing rate.

**Figure 5.**
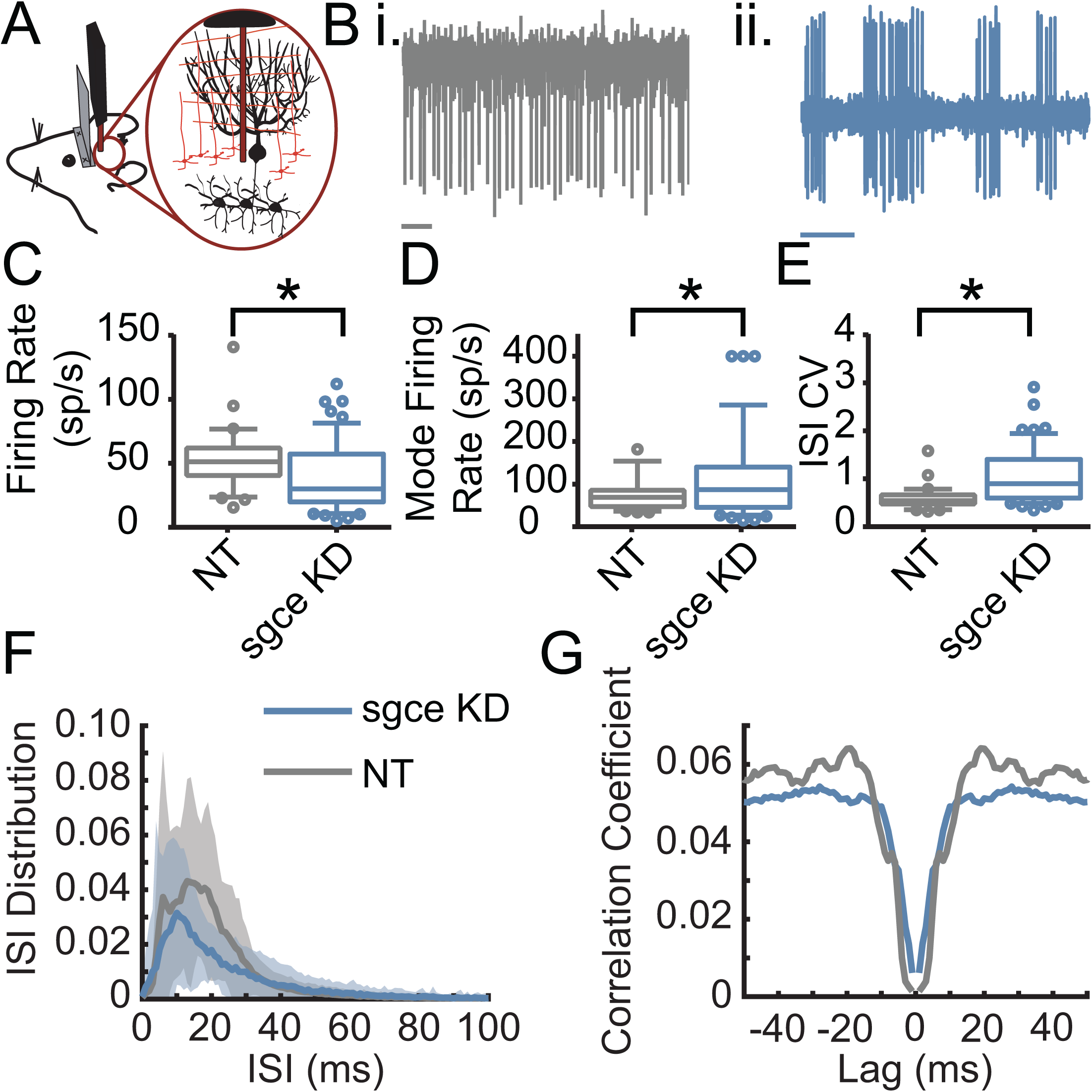
Purkinje cells fire aberrantly in sgce KD CB mice. (A) Experimental schematic. Extracellular electrophysiological recordings were made from Purkinje cells in awake, head-restrained mice. (B) Example traces of Purkinje cells from an NT CB (i) or sgce KD CB (ii) mouse. Scale bar represents 100 ms. (C) Average firing rates of Purkinje cells in NT CB and sgce KD CB animals (NT CB = 53.3 ± 24.1 spikes/s, n = 30, N = 4 and sgce KD CB = 39.2 ± 26.6 spikes/s, n = 57, N = 11, Mean ± S.D.; Welch’s t-test, p = 0.0028). (D) Mode firing rates of Purkinje cells in NT CB and sgce KD CB animals (NT CB = 77.5 ± 41.6 spikes/s and sgce KD CB = 115.0 ± 99.4 spikes/s, Mean ± S.D.; Welch’s t-test, p = 0.0158). (E) Interspike interval coefficients of variation of Purkinje cells in NT CB and sgce KD CB animals (NT CB = 0.60 ± 0.25 and sgce KD CB = 1.06 ± 0.57, Mean ± S.D.; Welch’s t-test, p < 0.0001). (F) Normalized ISI histogram of Purkinje cells in NT CB and sgce KD CB mice. (G) Autocorrelogram of Purkinje cells in NT CB and sgce KD CB mice.

## Discussion

The primary goal of this study was to test the hypothesis that acute knockdown of *Sgce,* the mouse homolog of the gene responsible for DYT11 in humans, in adult mice would more accurately model DYT11 by circumventing compensation that might occur in transgenic mouse models where the protein is absent throughout the development of the brain. We found that acute, shRNA-mediated knockdown of *Sgce* in the cerebellum, but not the basal ganglia, was sufficient to induce dystonia and other motor symptoms in adult mice. Consistent with what has been observed in patients, the motor symptoms of sgce KD CB mice were sensitive to alcohol, distinguishing this model from previous models of DYT11.

Previous genetic mouse models of DYT11 include both ubiquitous and cell-type specific *Sgce* knockout mice. Because the maternal *SGCE* allele is epigenetically silenced in humans, DYT11 is paternally inherited in the vast majority of cases. In mice, there is no expression of maternally-inherited *Sgce* (Yokoi et al., 2005). To reflect the inheritance pattern observed in humans, mouse models of DYT11 were generated by crossing male heterozygous *Sgce* knockout mice with wild-type females. Ubiquitous genetic knockout of *Sgce* resulted in subtle motor symptoms, including full body jerks and deficits in motor learning in a balance beam task (Yokoi et al., 2006). In contrast, sgce KD CB mice, which in most symptomatic cases have less than half of wild-type *Sgce* levels, exhibit myoclonic- and dystonic-like motor symptoms that are responsive to ethanol. Our success in generating symptomatic mice using an acute knockdown approach points to possible rodent-specific, developmental compensation for loss of *Sgce* in previous genetic mouse models of DYT11 (Xiao et al., 2017; Yokoi et al., 2006; Yokoi et al., 2012a; Yokoi et al., 2012b). Indeed, a recently-developed genetic model of DYT11 that used gene-trap technology to knock down the main and brain-specific isoforms of *Sgce* observed dystonic-like posturing between postnatal days 14 and 16, an important period of Purkinje cell development (Xiao et al., 2017). These symptoms did not persist into adulthood, however, supporting the hypothesis that developmental compensation may occur in genetic knockout animals. One possible source of compensation is the upregulation of genes with related functions. It has been shown that ε-SG forms complexes with the other members of the sarcoglycan family (α, β, γ,δ, and ζ), and also with members of the dystrophin-associated glycoprotein complex (DGC) in the brain (Waite et al., 2016). It is possible that upregulation of the other members in this complex during development could compensate for loss of *Sgce* in mice. The shRNA-mediated knockdown approach circumvents possible developmental compensation by knocking down the protein in the adult mouse and may be more effective than previous models of DYT11 for understanding how loss of *Sgce* leads to motor symptoms.

The model presented here is the first description of a mouse model of DYT11 that recapitulates the salient features of DYT11, namely, myoclonic-like movements and dystonia that were responsive to alcohol. However, in addition to overt dystonia, sgce KD CB mice also had difficulty ambulating normally and showed a mildly ataxic gate. While ataxia is uncommon in DYT11 patients, it has been observed in individuals with SGCE mutations (Drivenes et al., 2015; Sun et al., 2016). It is also worth noting that the most obvious motor symptoms in sgce KD CB mice were observed in the hind limbs and tails of the animals. While lower limb involvement has been reported in some DYT11 cases (Kobylecki et al., 2014), involvement of the trunk and upper extremities is far more common (Asmus et al., 2002). It is possible that our method of evaluating symptom severity based on activity in the open field biased us towards highlighting motor dysfunction in the hind limbs and the tail, and limited our ability to detect more severe motor symptoms in the forelimbs and trunk of sgce KD CB mice that can be more carefully scrutinized in a skilled forelimb reaching task. Lastly, sgce KD CB mice exhibited spinning when suspended by the tail. While there is no corresponding human symptom, understanding how loss of *Sgce* contributes to this symptom and how alcohol improves it may yield important insights into *Sgce* function.

The acute knockdown strategy further enables us to examine which parts of the brain are responsible for myoclonus and dystonia in DYT11. Our findings implicate the cerebellum as a central structure contributing to the motor symptoms in this disorder, in agreement with findings in human patients (Beukers et al., 2010; Carbon et al., 2013; Nitschke et al., 2006; van der Meer et al., 2012; van der Salm et al., 2013) and consistent with what has been reported in multiple dystonic rodent models (Campbell and Hess, 1998; Campbell et al., 1999; LeDoux et al., 1993; LeDoux et al., 1995; Neychev et al., 2008). We found that knockdown of *Sgce* specifically in the cerebellum led to dystonic-like movements that were responsive to alcohol. This suggests that the motor symptoms are a consequence of cerebellar dysfunction, supported by the aberrant activity recorded from DCN neurons and Purkinje cells, and that alcohol may act through the cerebellum to ameliorate them. We found that both Purkinje cells and DCN neurons exhibited irregular firing rates *in vivo*, indicated by an increased ISICV. These findings are consistent with our observations in many animal models of movement disorders, including ataxia and dystonia. In general, the more symptomatic the animal, the more irregular the cerebellar activity. In these animals, we found that irregular activity of Purkinje cells and DCN neurons was correlated with the presence of dystonia (Fremont et al., 2014; Fremont et al., 2017), in agreement with what has been reported in other rodent models of dystonia (Fremont et al., 2014; Fremont et al., 2015; Isaksen et al., 2017; LeDoux et al., 1998; LeDoux and Lorden, 1998, 2002; White and Sillitoe, 2017). While sgce KD CB mice exhibit dystonia, we did not perform EMG recordings in agonist and antagonist muscles at the time of recording. We were thus unable to determine whether highly irregular firing was time-locked to dystonic episodes. The increased ISI CV in sgce KD CB mice thus reflects a combination of acute dystonic, and persistent abnormal motor activity in these animals.

Cerebellar contribution to DYT11 was examined previously in a Purkinje cell-specific knockout of *Sgce.* In contrast to our findings and what was reported in ubiquitous *Sgce* null mice, the symptoms of Purkinje cell knockout mice were quite mild; they exhibited a small deficit in motor learning but had no robust motor abnormalities or myoclonic-like movements, suggesting brain regions other than the cerebellum might contribute to these motor symptoms (Yokoi et al., 2012a). However, one major difference between the present work and previous work is that our knockdown is not specific to Purkinje cells. It is possible that *Sgce*, which is expressed in DCN neurons in addition to Purkinje cells (Ritz et al., 2011), is important for the function of cerebellar output neurons. This is consistent with our finding that the average firing rate is decreased and the ISICV is increased in DCN neurons in sgce KD CB mice. The effect of *Sgce* knockdown in DCN neurons may provide an alternative explanation for the more severe motor phenotype of *Sgce* null mice compared to the Purkinje cell-specific knockout. Further experiments are required to determine whether it is the intrinsic activity of DCN neurons that contributes to motor symptoms, or whether it is their synaptic inputs, including Purkinje cells, which were also affected in our study, that lead to erratic cerebellar output and subsequent motor abnormalities. The symptomatic mouse model produced by acute knockdown of *Sgce* provides a unique opportunity to address these questions.

Because so much is known about the microcircuitry of the cerebellum, as well as the channels that contribute to the intrinsic firing rate of cerebellar neurons, understanding how the electrical properties of cerebellar neurons change in sgce KD CB mice will provide insight into the function of ε-SG under normal conditions. Understanding how alcohol acts to relieve dystonic symptoms would identify therapeutic targets for DYT11 in the cerebellum and help elucidate the mechanisms by which alcohol influences cerebellar output. A number of potential targets of ethanol in the brain have been described (Harris et al., 2008; Narahashi et al., 2001). One well-described and intensely-debated target of ethanol is the delta subunit-containing extrasynaptic GABA_A_ receptor (Hanchar et al., 2005; Wallner et al., 2003), which is responsible for maintaining tonic inhibitory current in cerebellar granule cells (Stell et al., 2003). Modulation of inhibitory currents by ethanol via extrasynaptic GABA_A_ receptors is a potential mechanism for restoring the firing rate and regularity of cerebellar neurons, thereby relieving dystonia.

While our findings are consistent with a growing body of evidence in support of a role for the cerebellum in DYT11, our results do not rule out the possibility that other brain regions also contribute to the symptoms observed in these mice. Knockdown of *Sgce* in the basal ganglia induced subtle motor defects in the mice. Frequently, sgce KD BG mice exhibited increased kinetic behavior specifically in the periphery of the open field chamber. An intriguing possibility is that this behavior is related more to the non-motor symptoms associated with DYT11, including obsessive compulsive disorder and anxiety.

It is now well-established that there are a number of direct connections between the cerebellum and the basal ganglia. Given the extensive connections between these brain regions and the contributions of both structures to motor control, it has been suggested that movement disorders should be considered less as disorders of a particular brain region, and more as disorders of the motor circuit. This is because dysfunction in one node, i.e. the cerebellum or basal ganglia, can influence the activity of other nodes (Bostan and Strick, 2018). With that in mind, two things become very clear. First, it is important from a basic science perspective to understand the site of initial dysfunction in the different dystonias in order to truly understand how abnormal movement results from a genetic mutation. In the case of DYT11, our approach suggests that loss-of-function of *Sgce* can lead to irregular activity of the cerebellum, resulting in motor symptoms. Understanding this link between genetic mutation and motor symptoms will yield important information about cerebellar function in health and disease. Second, the result of the interconnectivity of different brain structures is that there are multiple sites for therapeutic intervention. It has been shown that deep brain stimulation of the GPi, as well as the ventral intermediate nucleus of the thalamus (VIM) can improve symptoms in most DYT11 patients, with or without *SGCE* mutations (Azoulay-Zyss et al., 2011; Fernandez-Pajarin et al., 2016; Gruber et al., 2010; Kosutzka et al., 2018; Rocha et al., 2016; Rughani and Lozano, 2013). It is possible that DBS of the GPi or VIM may influence cerebellar-recipient regions of the thalamus, disrupting the irregular output from the cerebellum and, in this way, improving motor symptoms. Indeed, there is considerable overlap of cerebellar and pallidal projections in the thalamus (Hintzen et al., 2018). It is also possible, for example, that abnormal cerebellar output alters basal ganglia output and contributes to motor symptoms in dystonia. DBS of the GPi may disrupt the abnormal output from the basal ganglia, caused in part by cerebellar dysfunction, and restore some functionality to the system.

Taken together, our studies show that loss of function of *Sgce* in the cerebellum results in aberrant cerebellar output which in turn causes motor dysfunction, including jerky, myoclonic-like movements and dystonia, suggesting that the cerebellum may be a major of site of dysfunction in DYT11. The efficacy of ethanol in reducing the severity of the motor symptoms in sgce KD CB mice, as it does in DYT11 patients, further validates the model. Thus, the acute shRNA-mediated knockdown model of DYT11 described here not only provides a platform for further scrutiny of the mechanisms by which loss of ε-SG causes abnormal neuronal activity and motor dysfunction in DYT11, but also provides an opportunity to study how alcohol exerts its beneficial effects and, subsequently, identify alternative therapeutic strategies that mimic the effects of alcohol without its addictive and neurodegenerative consequences.

## Acknowledgements

We would like to acknowledge the scorers of the open field videos for all behavioral experiments, as well as our funding source, the National Institute of Neurological Disorders and Stroke (NINDS) of the NIH. We would further like to acknowledge that the affinity-purified antibody against ε-SG was a kind gift from Dr. Kevin Campbell (HHMI, University of Iowa).

## Methods

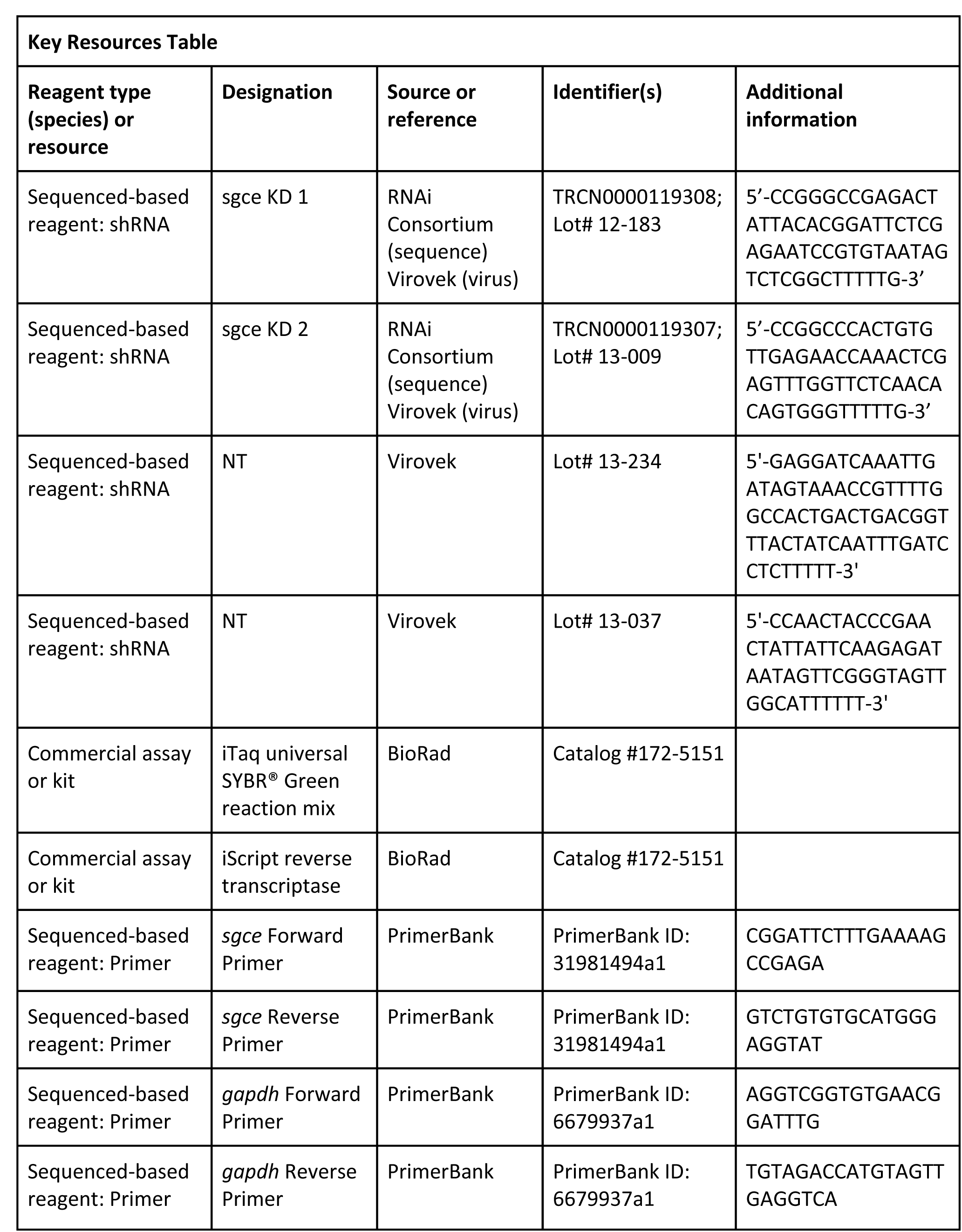
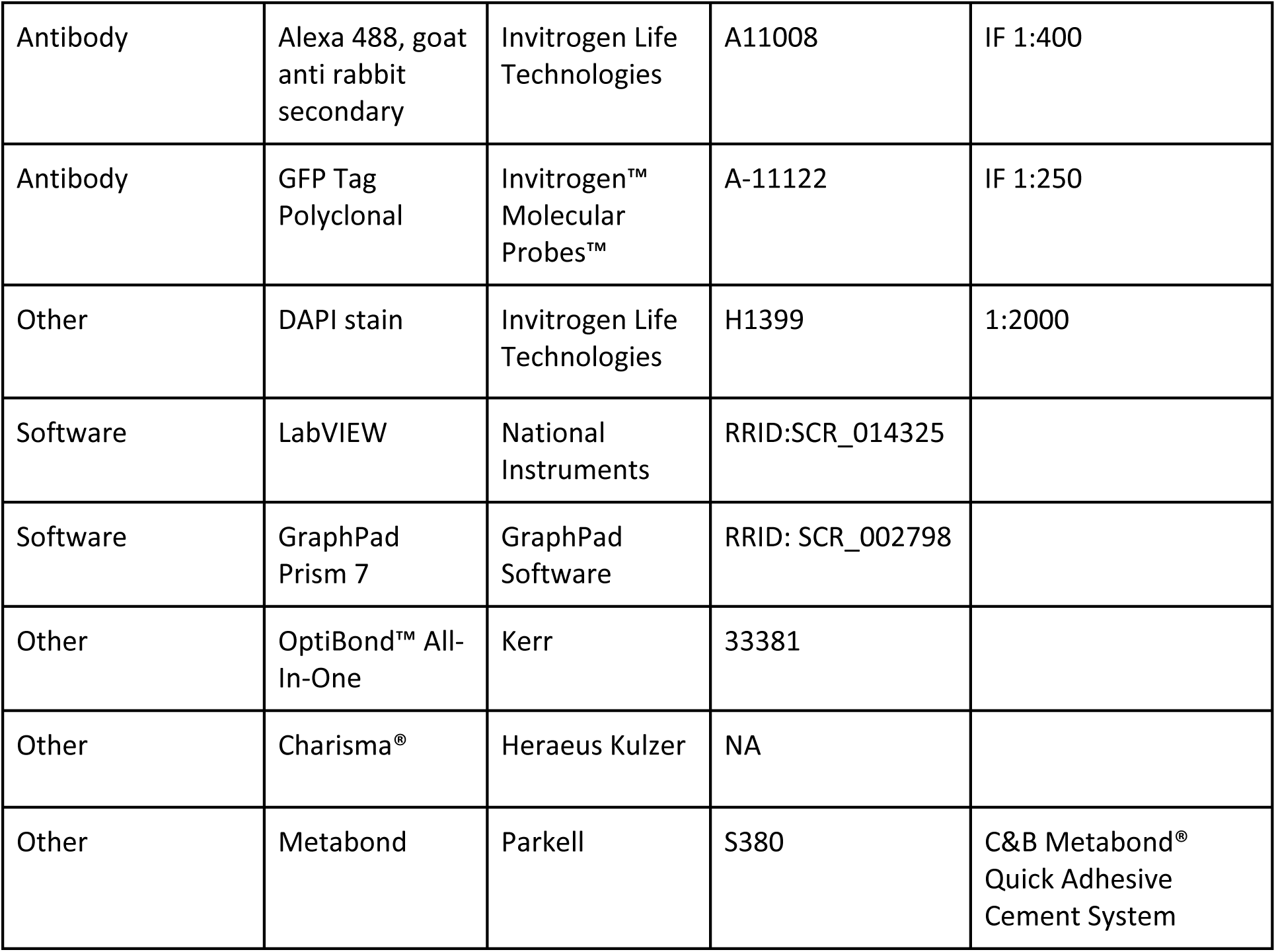

### Method details

Experiments were performed male and female C57BL/6 mice at least 6 weeks old in accordance with the guidelines set by Albert Einstein College of Medicine.

### shRNA Sequences

Two different shRNAs against unique parts of the *sgce* sequence generated originally by the RNAi consortium were identified and used in the following experiments (Key Resources Table). AAV9 compatible plasmids and AAV9 virus containing each shRNA were generated commercially (Virovek, Hayward, CA, AAV9-U6-shRNA119308-CMV-GFP (Lot# 12-183); AAV9-U6-shRNA_SGCE-CMV-GFP (Lot# 13-009), respectively). Each virus has an average titer of 2 x10^13^ vg/ml. A 0.22 µm filter sterilized solution containing the virus in DPBS buffer with 0.001% pluronic F-68 was directly injected into the cerebellum or basal ganglia of adult mice. Control AAV9 containing non-targeted (NT) shRNAs under the same promoter containing the same CMV driven GFP at equivalent titer were also purchased (Key Resources Table).

### Stereotaxic injection of shRNA

Male and female C57/Bl6 mice at least 6 weeks of age were anesthetized with 4% isoflurane and placed on a stereotaxic frame (David Kopf Instruments, Tujunga, CA). Isoflurane anesthesia was maintained at 1.5-2% throughout the course of the surgery, which was sufficient to prevent the animal from responding to a toe or tail pinch. A midline incision was made and the surface of the skull was cleaned of all connective tissue. Craniotomies were made over the cerebellum or basal ganglia, and 2 µl shRNA (AAV-SGCEshRNA-GFP or AAV-shNT-GFP) was injected into each injection site at a rate of 0.15 µl/min. After each injection, the syringe was retracted 50 µm and left for at least 5 minutes. For the cerebellum, the following coordinates were used: -6.00 AP, 0 ML, -1.5 DV and -6.96 AP, 0 ML, and 1.5 DV (cerebellar cortex) and -6.00 AP, ±1.8 ML, -2.3 DV (cerebellar nuclei). For the basal ganglia, the following coordinates were used: +0.5 AP, ±2.0 ML, -2.5 DV (striatum) and -0.5 AP, ±2.5 ML, -3.5 DV (GPi). A total of 8 µl was injected into the brain. In these experiments, every effort was made to include all injected animals. However, on rare occasions, an animal was excluded if post-hoc histology revealed a lack of expression of the virus in the targeted brain area or expression of the virus outside the targeted brain area.

### qRT-PCR

Mice were anesthetized with isoflurane until breathing slowed to 1 breath per 3 seconds and the animal was completely unresponsive to tail- or toe pinch. The animal was then quickly decapitated. The cerebellum was rapidly dissected, placed in a 1.5 mL Eppindorf tube, and frozen in liquid nitrogen.

Samples were then transported on dry ice to the Molecular Cytogenetic Core at Albert Einstein College of Medicine, where RNA extraction was performed. qRT-PCR was carried out on the CFX96 Touch™ Real-Time PCR Detection System (BioRad) with the iTaq™ Universal SYBR ® Green One-Step Kit (BioRad, Catalog# 172-5151). In total, RNA from the cerebella of 6 wild-type, 5 NT, 13 *sgce*KD1, and 7 *sgce*KD2 mice were examined. Primer sequences for *sgce* and *gapdh* were identified through PrimerBank (Spandidos et al., 2008; Spandidos et al., 2010; Wang and Seed, 2003), and oligonucleotides were commercially generated (Eurofins MWG Operon Oligos Tool, Thermo Fisher Scientific). Each 10 µl reaction contained 300nM each of forward and reverse primer and 100 ng RNA. All reactions were performed in duplicate or triplicate.

Analysis was performed using the 2^(-ΔΔC_T_) method (Livak and Schmittgen, 2001). In order to be used for analysis, the threshold cycle (C_T_) of two replicates had to be within 0.4 cycles of each other (standard deviation ≤ 0.283). The median C_T_ for each group of replicates was used. ΔC_T_ was calculated by subtracting the C_T_ for *gapdh*, a housekeeping gene control, from the C_T_ for *sgce* for each RNA sample. ΔΔC_T_ was calculated by subtracting the mean C_T_ for all WT samples from the C_T_ for each sample. Relative RNA quantity was then determined by calculating 2^(-ΔΔC_T_).

### Immunohistochemistry

Immunohistochemistry of GFP was used to confirm expression of the virus, which encodes GFP, in the targeted brain region. On rare occasions, animals were excluded when there was either no clear GFP expression in the target area or when there was GFP expressed in regions outside of the target area. Mice were anesthetized with isoflurane and transcardially perfused with phosphate buffered saline (PBS, Thermo Fisher Scientific, Waltham, MA) followed by 4% paraformaldehyde (PFA, Acros Organics, Thermo Fisher Scientific). The brains were dissected and fixed overnight in 4% PFA at 4°C. They were then rehydrated with 30% sucrose for 24-48 hours, rapidly frozen in Optimal Cutting Temperature Compound (OCT, Tissue-Tek) on dry ice, and stored at -80°C. Brains were cut on a cryostat (Leica CM3050 S) into 30 µm sections. Sections were stained with primary antibody against GFP (rabbit, 1:250, Molecular probes, A11122), followed by secondary antibody Alexa 488 (1:400, goat anti rabbit, Invitrogen Life Technologies A11008). Nuclei were labeled with DAPI (1:2000, Hoescht 33342, Invitrogen Life Technologies H1399). Images of sections were captured under a standard fluorescent microscope (Zeiss Axioskop 2 Plus).

### Dystonia Scale

The presence and severity of dystonia was quantified using a previously published scale (Calderon et al., 2011). Briefly, 0 = normal behavior; 1 = abnormal motor behavior, no dystonic postures; 2 = mild motor impairment, dystonic-like postures when disturbed; 3 = moderate impairment, frequent spontaneous dystonic postures; 4 = severe impairment, sustained dystonic postures. The videos were assessed independently by four observers who were blinded to the animal’s condition. The observers were trained with a video set containing representative examples for each score in which key characteristics were highlighted.

### Disability scale

In order to more accurately examine the range of motor symptoms observed in SGCE KD CB mice and measure the response of motor symptoms to ethanol, we generated the Disability Scale, which considers the frequency of both sustained dystonic-like postures and repetitive movements, and the extent of motor impairment. Behavior in the open field was scored by 4 colleagues blind to the condition of the animal as follows: 0 = Normal, the animal has no observable motor deficit; 1 = slight motor disability, the animal may exhibit unsteady gait or uncoordinated movement, but does not have any repetitive movements or sustained postures; 2 = Mild motor disability, ambulation is mildly impaired, the animal exhibits wide stance, very unsteady gait, back-walking, and/or occasional repetitive movements or sustained postures; 3 = Moderate motor disability, ambulation is moderately impaired; the animal does not properly ambulate, drags itself on its side, and may exhibit frequent sustained dystonic-like postures or repetitive movements; and 4 = severe motor disability, the animal is unable to properly ambulate for most of the duration of the observation, exhibits frequent repetitive movements, and/or sustained dystonic-like postures

### Spinning Scale

Sgce KD CB mice exhibited abnormal spinning when suspended by the tail. To quantify this spinning behavior, short video clips of the animal suspended by the tail were shown to 4 observers blind to the condition of the animal and scored as follows: 0 = Not present, the animal does not spin; 1 = mild, the animal spins but not rapidly and only for parts of the video; 2 = moderate, the animal spins rapidly, but not for the full duration of the video; and 3 = severe, the animal is spinning rapidly for the full duration of the video (Table 2, Video 5).

### Quantitative Open Field Analysis

Videos of equal length were analyzed using EthoVision XT 14, a commercially available tracking software program (Noldus).

### Ethanol and saline injections

A working solution of 0.2 g/mL ethanol (EtOH) in saline was prepared. After a 5 minute baseline period in the open field, sgce KD CB animals were subcutaneously injected with ethanol at a volume of 200 µl for a 20 g mouse, for a final dose of 2 g/kg EtOH. During control trials, animals were injected with an equal volume of physiological saline. The behavior in the open field was recorded every 15 minutes for 90 minutes following EtOH injection. Tor1a KD CB mice were generated exactly as previously described (Fremont et al., 2017). These animals were given the same ethanol treatment, and their behavior in the open field was recorded every 15 minutes for 60 minutes following EtOH injection. Saline-injected animals were followed up to 60 minutes following the injection.

### In vivo electrophysiology

An L-shaped or flat titanium bracket was affixed to the skull of sgce KD CB or NT CB mice with Charisma® (Heraeus Kulzer) or Metabond (C&B Metabond® Quick Adhesive Cement System, Parkell, S380) and dental cement. Craniotomies approximately 500 µm in diameter were made over the cerebellum for neuronal recordings. Recording sites were covered with a silicone adhesive (KWIK-SIL, WPI). Dental cement was used to construct a well around the recording site in order to hold saline during recording sessions.

Extracellular electrophysiological recordings were made from well-isolated single units using a tungsten electrode (Thomas Recording, 2-3 MΩ), which was advanced into the cerebellum until either the Purkinje cell layer or the DCN was reached. Purkinje cells were identified by location, firing rate, and the presence of complex spikes. DCN neurons were identified primarily by location and firing rate. Neurophysiological signals were amplified 2000x using a custom built amplifier and digitized at 20 kHz using a National Instruments BNC-2110. Waveforms were sorted offline by principal component analysis (Plexon).

### Statistics

GraphPad Prism 7 (GraphPad Software) was used to perform all statistical analyses. Data were assessed for normality using Shapiro-Wilk normality test. Non-normal data were compared using a non-parametric Wilcoxon test. Between group analysis was performed using the Holm-Sidak method. Each row was analyzed individually, without assuming a consistent SD. Normally distributed data sets were statistically compared based on the two-tailed, paired Student’s t-test or one-way ANOVA with Tukey’s correction for multiple comparisons. Data are reported in text as mean ± S.D. unless otherwise stated.

---

**Figure 2 – Figure Supplement 1.**
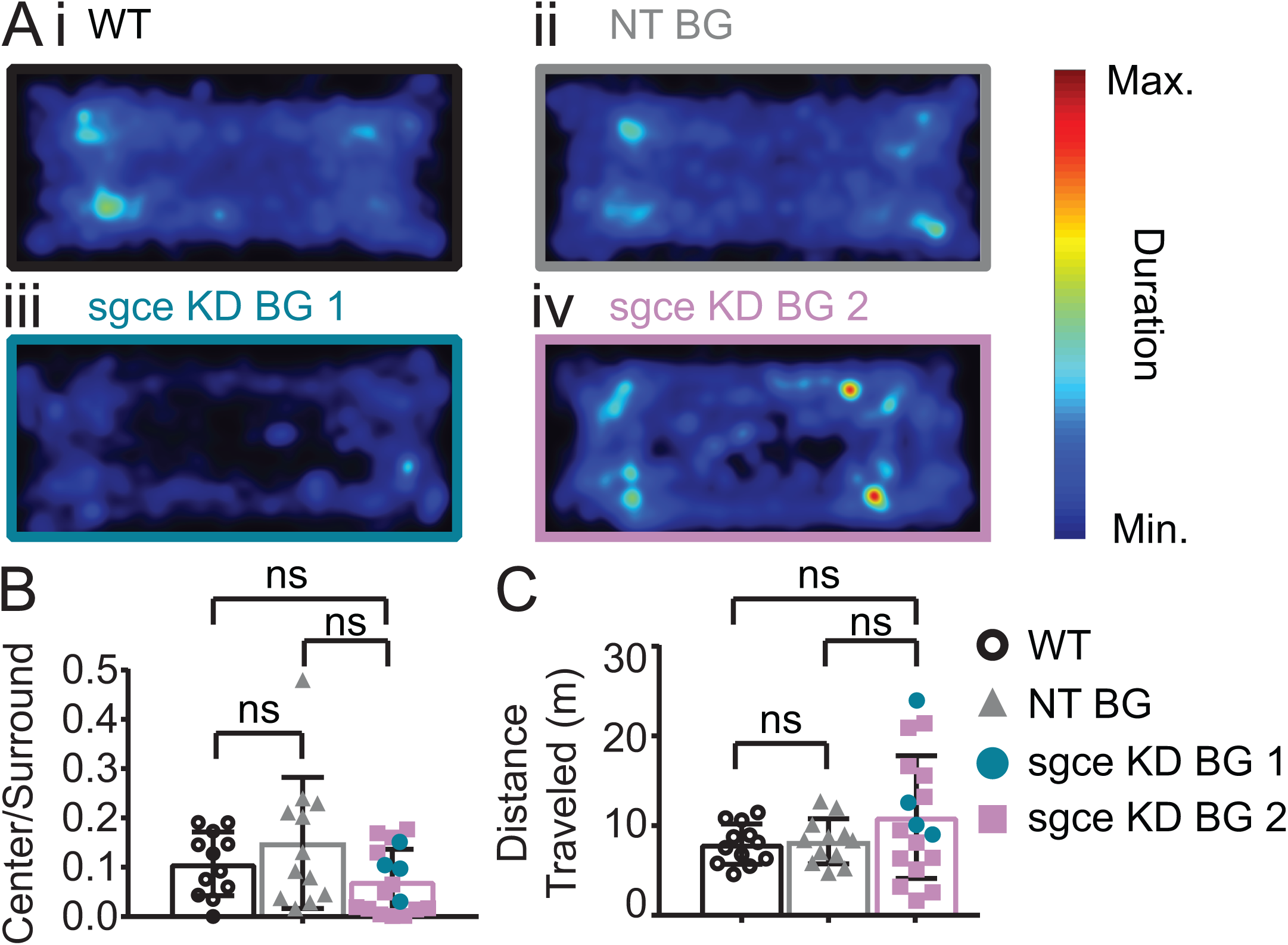
Sgce KD BG mice spend qualitatively less time in the center of the open field with no significant change in the ratio of distance traveled. (A) Average duration spent per pixel in the open field chamber (WT: N = 12, NT BG: N = 12, sgce KD BG 1: N = 4; sgce KD BG 2: N = 13). (B) Ratio of duration of time spent in the center, to the duration of time spent in the surround in a 5 minute video (Mean + S.D., WT: N = 12, NT BG: N = 12, sgce KD BG 1: N = 4; sgce KD BG 2: N = 13, 1way ANOVA, Holm-Sidak’s multiple comparisons test, p > 0.05 for all comparisons). (C) Distance traveled (m) in a 5 minute video (Mean + S.D., WT: N = 12, NT BG: N = 12, sgce KD BG 1: N = 4; sgce KD BG 2: N = 13, 1way ANOVA, Holm-Sidak’s multiple comparisons test, p > 0.05 for all comparisons).

**Figure 2 – Figure Supplement 2.**
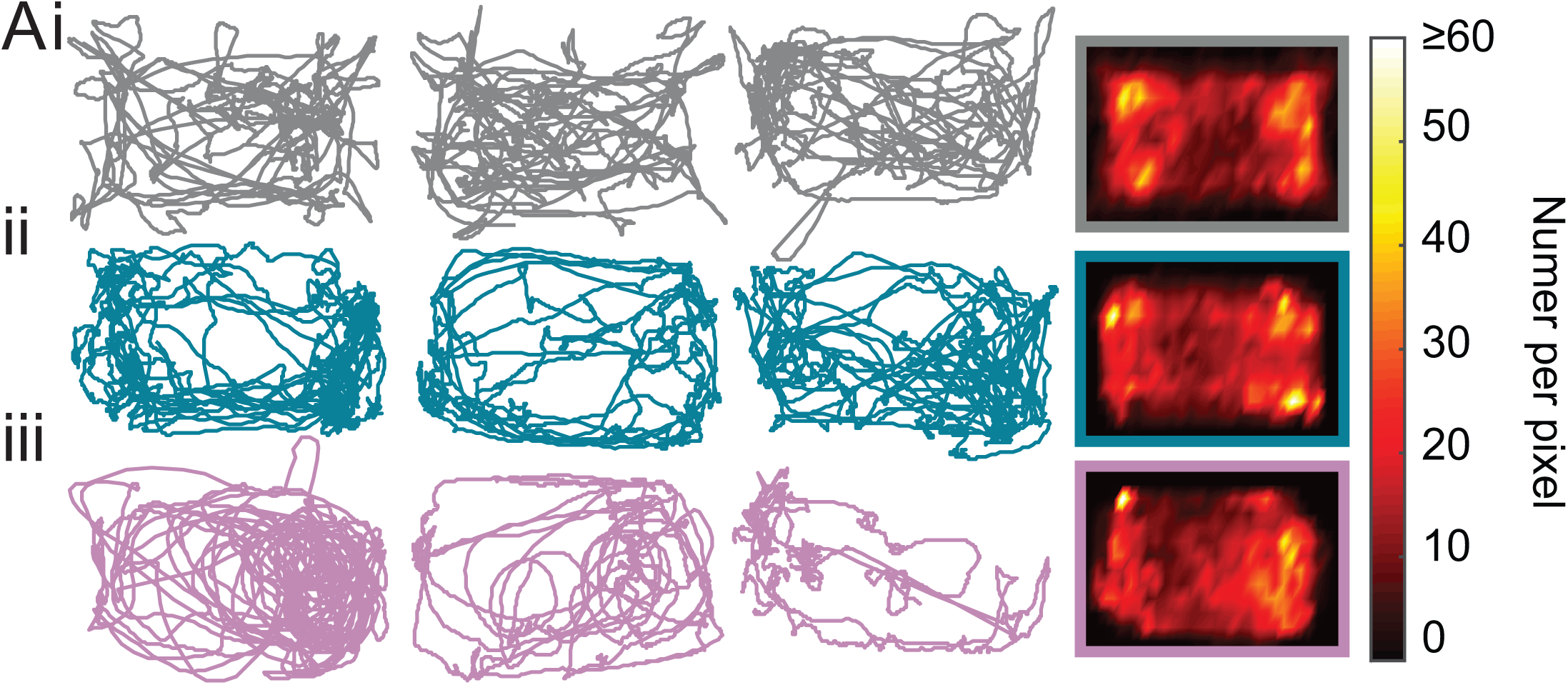
Sgce KD CB mice did not show a preference for the periphery and avoidance of the center. (A) sgce KD CB 1 (ii) and sgce KD CB 2 mice (iii) showed no difference in ambulation in the open field chamber compared to NT CB (i) mice. Columns 1-3 are example tracks from individual mice, and column 4 is the average (NT CB: N = 14, sgce KD CB 1: N = 11, sgce KD CB 2: N = 9).

**Figure 3 – Figure Supplement 1.**
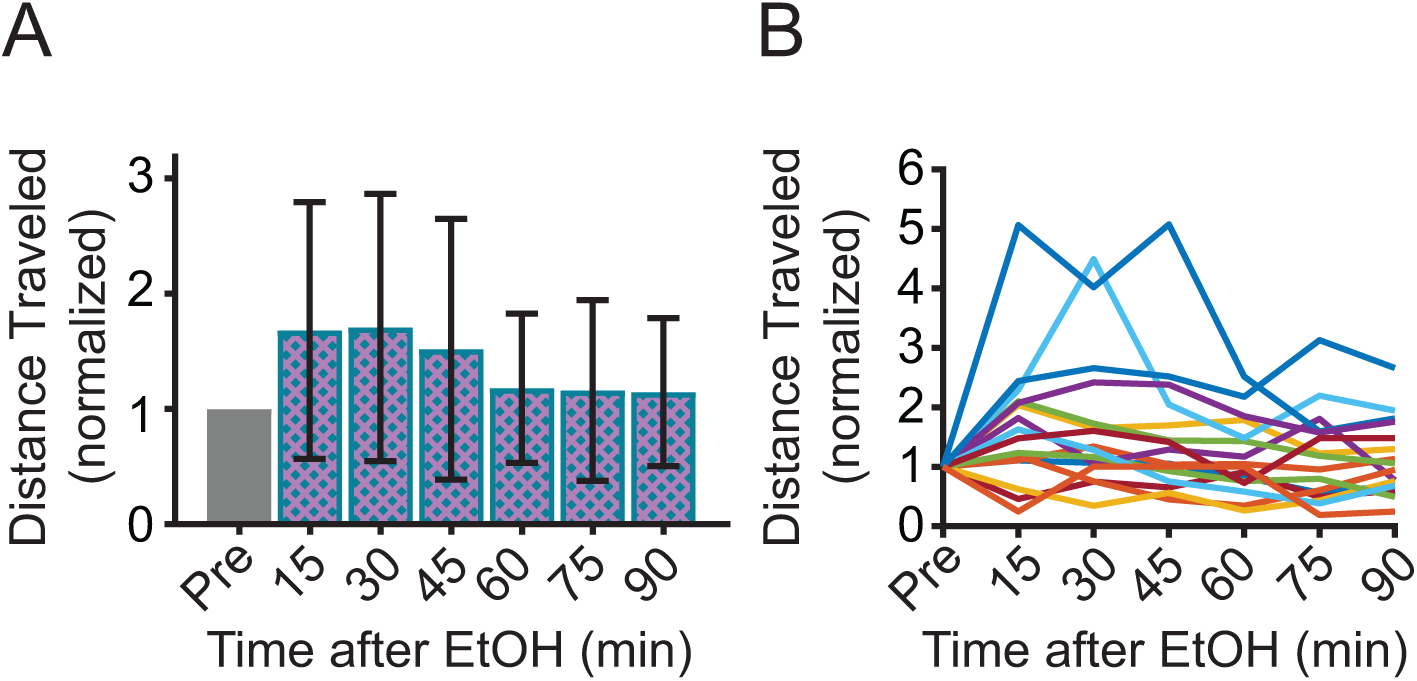
The effect of EtOH on distance travelled varies among individual mice. (A) Total distance traveled in the open field after ethanol injection, normalized to distance traveled before ethanol (Pre) (1way ANOVA, p = 0.0148, N = 16, Dunnett’s multiple comparisson’ test, p> 0.05 at each time point). (B) Data in (A) presented for each mouse.

**Figure 3 – Supplement 2.**
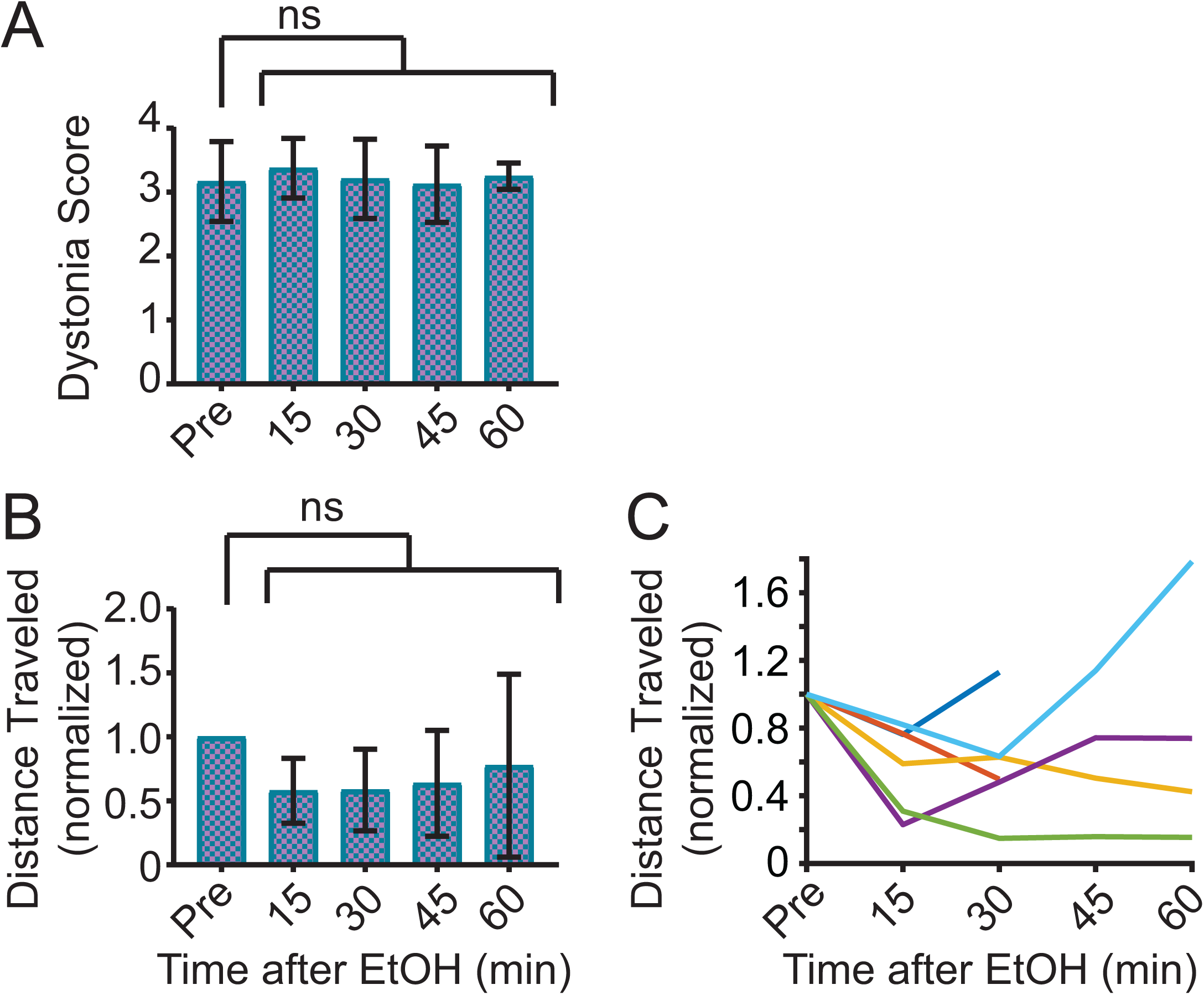
Saline does not improve symptoms in sgce KD CB mice. (A) Dystonia score of sgce KD CB mice after saline injection. Saline had no effect on the dystonia score of sgce KD CB mice (p = 0.9516, 1wayANVOA, N = 6). (B) Total distance traveled in the open field after EtOH injection, normalized to distance traveled before ethanol (Pre) (p = 0.2851, 1way ANOVA, N = 6). (C) Data in (B) presented for each mouse.

## Video Legends

**Video1.** sgce KD CB 1 and sgce KD CB 2 mice have dystonia, as evidenced by a dystonia score of greater than or equal to 2 in the open field, while NT CB mice do not develop dystonia (p < 0.01, Wilcoxon matched-pairs signed rank test, sgce KD CB 1: N = 39; sgce KD CB 2: N = 40; NT CB: N = 16).

**Video2.** In addition to dystonia, sgce KD CB mice exhibit myoclonic-like movements in the open field and spin when suspended by the tail.

**Video3.** Neither sgce KD BG nor NT BG mice developed dystonia, as measured by a score of greater than or equal to 2 in the open field.

**Video4.** EtOH relieves motor symptoms of sgce KD CB mice, as measured on the Disability Scale (p < 0.0001, 1way ANOVA, N = 16), Spinning Scale Scale (, p < 0.0001, 1way ANOVA, N = 19), and Dystonia Scale (p < 0.0001, 1way ANOVA, N = 16), but it does not relieve dystonic symptoms of mice injected with shRNA against tor1a, an acute shRNA knockdown model of DYT1 (p = 0.2391, 1way ANOVA, N = 5).

**Video5.** Saline had no effect on the motor symptoms of sgce KD CB mice (p = 0.9517, 1way ANOVA, N = 6).

## References

1. Albanese, A., Bhatia, K., Bressman, S.B., Delong, M.R., Fahn, S., Fung, V.S., Hallett, M., Jankovic, J., Jinnah, H.A., Klein, C., et al. (2013). Phenomenology and classification of dystonia: A consensus update. Movement disorders: official journal of the Movement Disorder Society 28, 863–873.

2. Asmus, F., Zimprich, A., Tezenas Du Montcel, S., Kabus, C., Deuschl, G., Kupsch, A., Ziemann, U., Castro, M., Kuhn, A.A., Strom, T.M., et al. (2002). Myoclonus-dystonia syndrome: epsilon-sarcoglycan mutations and phenotype. Annals of neurology 52, 489–492.

3. Azoulay-Zyss, J., Roze, E., Welter, M.L., Navarro, S., Yelnik, J., Clot, F., Bardinet, E., Karachi, C., Dormont, D., Galanaud, D., et al. (2011). Bilateral deep brain stimulation of the pallidum for myoclonus-dystonia due to epsilon-sarcoglycan mutations: a pilot study. Archives of neurology 68, 94–98.

4. Belmeguenai, A., Botta, P., Weber, J.T., Carta, M., De Ruiter, M., De Zeeuw, C.I., Valenzuela, C.F., and Hansel, C. (2008). Alcohol impairs long-term depression at the cerebellar parallel fiber-Purkinje cell synapse. Journal of neurophysiology 100, 3167–3174.

5. Beukers, R.J., Foncke, E.M., van der Meer, J.N., Nederveen, A.J., de Ruiter, M.B., Bour, L.J., Veltman, D.J., and Tijssen, M.A. (2010). Disorganized sensorimotor integration in mutation-positive myoclonus-dystonia: a functional magnetic resonance imaging study. Archives of neurology 67, 469–474.

6. Beukers, R.J., van der Meer, J.N., van der Salm, S.M., Foncke, E.M., Veltman, D.J., and Tijssen, M.A. (2011). Severity of dystonia is correlated with putaminal gray matter changes in myoclonus-dystonia. European journal of neurology: the official journal of the European Federation of Neurological Societies 18, 906–912.

7. Bostan, A.C., and Strick, P.L. (2018). The basal ganglia and the cerebellum: nodes in an integrated network. Nature reviews Neuroscience 19, 338–350.

8. Calderon, D.P., Fremont, R., Kraenzlin, F., and Khodakhah, K. (2011). The neural substrates of rapid-onset Dystonia-Parkinsonism. Nat Neurosci 14, 357–365.

9. Campbell, D.B., and Hess, E.J. (1998). Cerebellar circuitry is activated during convulsive episodes in the tottering (tg/tg) mutant mouse. Neuroscience 85, 773–783.

10. Campbell, D.B., North, J.B., and Hess, E.J. (1999). Tottering mouse motor dysfunction is abolished on the Purkinje cell degeneration (pcd) mutant background. ExpNeurol 160, 268–278.

11. Carbon, M., Raymond, D., Ozelius, L., Saunders-Pullman, R., Frucht, S., Dhawan, V., Bressman, S., and Eidelberg, D. (2013). Metabolic changes in DYT11 myoclonus-dystonia. Neurology 80, 385–391.

12. Carta, M., Mameli, M., and Valenzuela, C.F. (2004). Alcohol enhances GABAergic transmission to cerebellar granule cells via an increase in Golgi cell excitability. The Journal of neuroscience: the official journal of the Society for Neuroscience 24, 3746–3751.

13. Charlesworth, G., Bhatia, K.P., and Wood, N.W. (2013). The genetics of dystonia: new twists in an old tale. Brain: a journal of neurology 136, 2017–2037.

14. Chu, N.S. (1983). Effects of ethanol on rat cerebellar Purkinje cells. The International journal of neuroscience 21, 265–277.

15. Draganski, B., Thun-Hohenstein, C., Bogdahn, U., Winkler, J., and May, A. (2003). “Motor circuit” gray matter changes in idiopathic cervical dystonia. Neurology 61, 1228–1231.

16. Dresel, C., Li, Y., Wilzeck, V., Castrop, F., Zimmer, C., and Haslinger, B. (2014). Multiple changes of functional connectivity between sensorimotor areas in focal hand dystonia. Journal of neurology, neurosurgery, and psychiatry 85, 1245–1252.

17. Drivenes, B., Born, A.P., Ek, J., Dunoe, M., and Uldall, P.V. (2015). A child with myoclonus-dystonia (DYT11) misdiagnosed as atypical opsoclonus myoclonus syndrome. Eur J Paediatr Neurol 19, 719–721.

18. Fernandez-Pajarin, G., Sesar, A., Relova, J.L., Ares, B., Jimenez-Martin, I., Blanco-Arias, P., Gelabert-Gonzalez, M., and Castro, A. (2016). Bilateral pallidal deep brain stimulation in myoclonus-dystonia: our experience in three cases and their follow-up. Acta Neurochir (Wien) 158, 2023–2028.

19. Fremont, R., Calderon, D.P., Maleki, S., and Khodakhah, K. (2014). Abnormal high-frequency burst firing of cerebellar neurons in rapid-onset dystonia-parkinsonism. J Neurosci 34, 11723–11732.

20. Fremont, R., and Khodakhah, K. (2012). Alternative approaches to modeling hereditary dystonias. Neurotherapeutics: the journal of the American Society for Experimental NeuroTherapeutics 9, 315–322.

21. Fremont, R., Tewari, A., Angueyra, C., and Khodakhah, K. (2017). A role for cerebellum in the hereditary dystonia DYT1. Elife 6.

22. Fremont, R., Tewari, A., and Khodakhah, K. (2015). Aberrant Purkinje cell activity is the cause of dystonia in a shRNA-based mouse model of Rapid Onset Dystonia-Parkinsonism. Neurobiol Dis 82, 200–212.

23. Grabowski, M., Zimprich, A., Lorenz-Depiereux, B., Kalscheuer, V., Asmus, F., Gasser, T., Meitinger, T., and Strom, T.M. (2003). The epsilon-sarcoglycan gene (SGCE), mutated in myoclonus-dystonia syndrome, is maternally imprinted. Eur J Hum Genet 11, 138–144.

24. Gruber, D., Kuhn, A.A., Schoenecker, T., Kivi, A., Trottenberg, T., Hoffmann, K.T., Gharabaghi, A., Kopp, U.A., Schneider, G.H., Klein, C., et al. (2010). Pallidal and thalamic deep brain stimulation in myoclonus-dystonia. Movement disorders: official journal of the Movement Disorder Society 25, 1733–1743.

25. Hanchar, H.J., Dodson, P.D., Olsen, R.W., Otis, T.S., and Wallner, M. (2005). Alcohol-induced motor impairment caused by increased extrasynaptic GABA(A) receptor activity. Nature neuroscience 8, 339–345.

26. Harris, R.A., Trudell, J.R., and Mihic, S.J. (2008). Ethanol’s molecular targets. Science signaling 1, re7.

27. He, Q., Titley, H., Grasselli, G., Piochon, C., and Hansel, C. (2013). Ethanol affects NMDA receptor signaling at climbing fiber-Purkinje cell synapses in mice and impairs cerebellar LTD. Journal of neurophysiology 109, 1333–1342.

28. Hendrix, C.M., and Vitek, J.L. (2012). Toward a network model of dystonia. Annals of the New York Academy of Sciences 1265, 46–55.

29. Hintzen, A., Pelzer, E.A., and Tittgemeyer, M. (2018). Thalamic interactions of cerebellum and basal ganglia. Brain Struct Funct 223, 569–587.

30. Hjermind, L.E., Vissing, J., Asmus, F., Krag, T., Lochmuller, H., Walter, M.C., Erdal, J., Blake, D.J., and Nielsen, J.E. (2008). No muscle involvement in myoclonus-dystonia caused by epsilon-sarcoglycan gene mutations. European journal of neurology: the official journal of the European Federation of Neurological Societies 15, 525–529.

31. Isaksen, T.J., Kros, L., Vedovato, N., Holm, T.H., Vitenzon, A., Gadsby, D.C., Khodakhah, K., and Lykke-Hartmann, K. (2017). Hypothermia-induced dystonia and abnormal cerebellar activity in a mouse model with a single disease-mutation in the sodium-potassium pump. PLoS Genet 13, e1006763.

32. Jinnah, H.A., and Hess, E.J. (2006). A new twist on the anatomy of dystonia: the basal ganglia and the cerebellum? Neurology 67, 1740–1741.

33. Kobylecki, C., Damodaran, D., Kerr, B., Newton, R.W., and Silverdale, M.A. (2014). Prominent Lower-Limb Involvement in a Family with Myoclonus-Dystonia. Mov Disord Clin Pract 1, 115–117.

34. Kosutzka, Z., Tisch, S., Bonnet, C., Ruiz, M., Hainque, E., Welter, M.L., Viallet, F., Karachi, C., Navarro, S., Jahanshahi, M., et al. (2018). Long-term GPi-DBS improves motor features in myoclonus-dystonia and enhances social adjustment. Movement disorders: official journal of the Movement Disorder Society.

35. Kyllerman, M., Forsgren, L., Sanner, G., Holmgren, G., Wahlstrom, J., and Drugge, U. (1990). Alcohol-responsive myoclonic dystonia in a large family: dominant inheritance and phenotypic variation. Movement disorders: official journal of the Movement Disorder Society 5, 270–279.

36. LeDoux, M.S., Hurst, D.C., and Lorden, J.F. (1998). Single-unit activity of cerebellar nuclear cells in the awake genetically dystonic rat. Neuroscience 86, 533–545.

37. LeDoux, M.S., and Lorden, J.F. (1998). Abnormal cerebellar output in the genetically dystonic rat. AdvNeurol 78, 63–78.

38. LeDoux, M.S., and Lorden, J.F. (2002). Abnormal spontaneous and harmaline-stimulated Purkinje cell activity in the awake genetically dystonic rat. ExpBrain Res 145, 457–467.

39. LeDoux, M.S., Lorden, J.F., and Ervin, J.M. (1993). Cerebellectomy eliminates the motor syndrome of the genetically dystonic rat. ExpNeurol 120, 302–310.

40. LeDoux, M.S., Lorden, J.F., and Meinzen-Derr, J. (1995). Selective elimination of cerebellar output in the genetically dystonic rat. Brain research 697, 91–103.

41. Livak, K.J., and Schmittgen, T.D. (2001). Analysis of relative gene expression data using real-time quantitative PCR and the 2(-Delta Delta C(T)) Method. Methods 25, 402–408.

42. Marelli, C., Canafoglia, L., Zibordi, F., Ciano, C., Visani, E., Zorzi, G., Garavaglia, B., Barzaghi, C., Albanese, A., Soliveri, P., et al. (2008). A neurophysiological study of myoclonus in patients with DYT11 myoclonus-dystonia syndrome. Movement disorders: official journal of the Movement Disorder Society 23, 2041–2048.

43. Narahashi, T., Kuriyama, K., Illes, P., Wirkner, K., Fischer, W., Muhlberg, K., Scheibler, P., Allgaier, C., Minami, K., Lovinger, D., et al. (2001). Neuroreceptors and ion channels as targets of alcohol. Alcoholism, clinical and experimental research 25, 182S–188S.

44. Nardocci, N., Zorzi, G., Barzaghi, C., Zibordi, F., Ciano, C., Ghezzi, D., and Garavaglia, B. (2008). Myoclonus-dystonia syndrome: clinical presentation, disease course, and genetic features in 11 families. Movement disorders: official journal of the Movement Disorder Society 23, 28–34.

45. Neychev, V.K., Fan, X., Mitev, V.I., Hess, E.J., and Jinnah, H.A. (2008). The basal ganglia and cerebellum interact in the expression of dystonic movement. Brain: a journal of neurology 131, 2499–2509.

46. Nitschke, M.F., Erdmann, C., Trillenberg, P., Sprenger, A., Kock, N., Sperner, J., and Klein, C. (2006). Functional MRI reveals activation of a subcortical network in a 5-year-old girl with genetically confirmed myoclonus-dystonia. Neuropediatrics 37, 79–82.

47. Peall, K.J., Waite, A.J., Blake, D.J., Owen, M.J., and Morris, H.R. (2011). Psychiatric disorders, myoclonus dystonia, and the epsilon-sarcoglycan gene: a systematic review. Movement disorders: official journal of the Movement Disorder Society 26, 1939–1942.

48. Prell, T., Peschel, T., Kohler, B., Bokemeyer, M.H., Dengler, R., Gunther, A., and Grosskreutz, J. (2013). Structural brain abnormalities in cervical dystonia. BMC Neurosci 14, 123.

49. Raymond, D., and Ozelius, L. (1993). Myoclonus-Dystonia. In GeneReviews((R)), M.P. Adam, H.H. Ardinger, R.A. Pagon, S.E. Wallace, L.J.H. Bean, K. Stephens, and A. Amemiya, eds. (Seattle (WA)).

50. Raymond, D., Saunders-Pullman, R., de Carvalho Aguiar, P., Schule, B., Kock, N., Friedman, J., Harris, J., Ford, B., Frucht, S., Heiman, G.A., et al. (2008). Phenotypic spectrum and sex effects in eleven myoclonus-dystonia families with epsilon-sarcoglycan mutations. Movement disorders: official journal of the Movement Disorder Society 23, 588–592.

51. Ritz, K., van Schaik, B.D., Jakobs, M.E., van Kampen, A.H., Aronica, E., Tijssen, M.A., and Baas, F. (2011). SGCE isoform characterization and expression in human brain: implications for myoclonus-dystonia pathogenesis? Eur J Hum Genet 19, 438–444.

52. Rocha, H., Linhares, P., Chamadoira, C., Rosas, M.J., and Vaz, R. (2016). Early deep brain stimulation in patients with myoclonus-dystonia syndrome. Journal of clinical neuroscience: official journal of the Neurosurgical Society of Australasia 27, 17–21.

53. Rogers, J., Siggins, G.R., Schulman, J.A., and Bloom, F.E. (1980). Physiological correlates of ethanol intoxication tolerance, and dependence in rat cerebellar Purkinje cells. Brain research 196, 183–198.

54. Roze, E., Saudou, F., and Caboche, J. (2008). Pathophysiology of Huntington’s disease: from huntingtin functions to potential treatments. Current opinion in neurology 21, 497–503.

55. Rughani, A.I., and Lozano, A.M. (2013). Surgical treatment of myoclonus dystonia syndrome. Movement disorders: official journal of the Movement Disorder Society 28, 282–287.

56. Shakkottai, V.G. (2014). Physiologic changes associated with cerebellar dystonia. Cerebellum 13, 637–644.

57. Sinclair, J.G., Lo, G.F., and Tien, A.F. (1980). The effects of ethanol on cerebellar Purkinje cells in naive and alcohol-dependent rats. Canadian journal of physiology and pharmacology 58, 429–432.

58. Song, C.H., Bernhard, D., Hess, E.J., and Jinnah, H.A. (2014). Subtle microstructural changes of the cerebellum in a knock-in mouse model of DYT1 dystonia. Neurobiol Dis 62, 372–380.

59. Spandidos, A., Wang, X., Wang, H., Dragnev, S., Thurber, T., and Seed, B. (2008). A comprehensive collection of experimentally validated primers for Polymerase Chain Reaction quantitation of murine transcript abundance. BMC Genomics 9, 633.

60. Spandidos, A., Wang, X., Wang, H., and Seed, B. (2010). PrimerBank: a resource of human and mouse PCR primer pairs for gene expression detection and quantification. Nucleic acids research 38, D792–799.

61. Steeves, T.D., Day, L., Dykeman, J., Jette, N., and Pringsheim, T. (2012). The prevalence of primary dystonia: a systematic review and meta-analysis. Movement disorders: official journal of the Movement Disorder Society 27, 1789–1796.

62. Stell, B.M., Brickley, S.G., Tang, C.Y., Farrant, M., and Mody, I. (2003). Neuroactive steroids reduce neuronal excitability by selectively enhancing tonic inhibition mediated by delta subunit-containing GABAA receptors. Proceedings of the National Academy of Sciences of the United States of America 100, 14439–14444.

63. Sun, L., Wu, J., Liu, C., and Lin, W. (2016). Report of progressive myoclonus ataxia (PMA) in two Chinese pedigrees. Neurol Res 38, 893–896.

64. van der Meer, J.N., Beukers, R.J., van der Salm, S.M., Caan, M.W., Tijssen, M.A., and Nederveen, A.J. (2012). White matter abnormalities in gene-positive myoclonus-dystonia. Movement disorders: official journal of the Movement Disorder Society 27, 1666–1672.

65. van der Salm, S.M., van der Meer, J.N., Nederveen, A.J., Veltman, D.J., van Rootselaar, A.F., and Tijssen, M.A. (2013). Functional MRI study of response inhibition in myoclonus dystonia. Experimental neurology 247, 623–629.

66. Waite, A.J., Carlisle, F.A., Chan, Y.M., and Blake, D.J. (2016). Myoclonus dystonia and muscular dystrophy: varepsilon-sarcoglycan is part of the dystrophin-associated protein complex in brain. Movement disorders: official journal of the Movement Disorder Society 31, 1694–1703.

67. Wallner, M., Hanchar, H.J., and Olsen, R.W. (2003). Ethanol enhances alpha 4 beta 3 delta and alpha 6 beta 3 delta gamma-aminobutyric acid type A receptors at low concentrations known to affect humans. Proceedings of the National Academy of Sciences of the United States of America 100, 15218–15223.

68. Wang, X., and Seed, B. (2003). A PCR primer bank for quantitative gene expression analysis. Nucleic acids research 31, e154.

69. Weissbach, A., Kasten, M., Grunewald, A., Bruggemann, N., Trillenberg, P., Klein, C., and Hagenah, J. (2013). Prominent psychiatric comorbidity in the dominantly inherited movement disorder myoclonus-dystonia. Parkinsonism Relat Disord 19, 422–425.

70. Welter, M.L., Grabli, D., Karachi, C., Jodoin, N., Fernandez-Vidal, S., Brun, Y., Navarro, S., Rogers, A., Cornu, P., Pidoux, B., et al. (2015). Pallidal activity in myoclonus dystonia correlates with motor signs. Movement disorders: official journal of the Movement Disorder Society 30, 992–996.

71. White, J.J., and Sillitoe, R.V. (2017). Genetic silencing of olivocerebellar synapses causes dystonia-like behaviour in mice. Nature communications 8, 14912.

72. Xiao, J., Vemula, S.R., Xue, Y., Khan, M.M., Carlisle, F.A., Waite, A.J., Blake, D.J., Dragatsis, I., Zhao, Y., and LeDoux, M.S. (2017). Role of major and brain-specific Sgce isoforms in the pathogenesis of myoclonus-dystonia syndrome. Neurobiol Dis 98, 52–65.

73. Yokoi, F., Dang, M.T., Li, J., and Li, Y. (2006). Myoclonus, motor deficits, alterations in emotional responses and monoamine metabolism in epsilon-sarcoglycan deficient mice. J Biochem 140, 141–146.

74. Yokoi, F., Dang, M.T., Mitsui, S., and Li, Y. (2005). Exclusive paternal expression and novel alternatively spliced variants of epsilon-sarcoglycan mRNA in mouse brain. FEBS Lett 579, 4822–4828.

75. Yokoi, F., Dang, M.T., Yang, G., Li, J., Doroodchi, A., Zhou, T., and Li, Y. (2012a). Abnormal nuclear envelope in the cerebellar Purkinje cells and impaired motor learning in DYT11 myoclonus-dystonia mouse models. Behav Brain Res 227, 12–20.

76. Yokoi, F., Dang, M.T., Zhou, T., and Li, Y. (2012b). Abnormal nuclear envelopes in the striatum and motor deficits in DYT11 myoclonus-dystonia mouse models. Human molecular genetics 21, 916–925.

77. Zimprich, A., Grabowski, M., Asmus, F., Naumann, M., Berg, D., Bertram, M., Scheidtmann, K., Kern, P., Winkelmann, J., Muller-Myhsok, B., et al. (2001). Mutations in the gene encoding epsilon-sarcoglycan cause myoclonus-dystonia syndrome. Nature genetics 29, 66–69.

